# Sirtuin 2 controls global protein synthesis by regulating Rheb-GTPase

**DOI:** 10.1101/2025.03.31.646364

**Authors:** Amarjeet Shrama, Yanlin Zi, Anwit Shriniwas Pandit, Kirtika Jha, Vikrant Kumar Sinha, Venkatraman Ravi, Souvik Ghosh, Dimple Nagesh, Bhoomika Shivanaiah, Danish Khan, Arathi Bangalore Prabhashankar, Thoniparambil Sunil Sumi, Sukanya Raghu, Anand Srivastava, Mahavir Singh, Hening Lin, Nagalingam R. Sundaresan

## Abstract

Upregulated global protein synthesis is associated with the development and progression of several diseases and disorders. Strategies like calorie restriction and pharmacological inhibition of protein synthesis, have exhibited health-promoting effects. However, the complex molecular events that regulate global protein synthesis are not completely understood. Here, we report that SIRT2, a histone deacylase, negatively regulates global protein synthesis by inhibiting the mTORC1 pathway via deacetylating Rheb and promoting Rheb degradation. Our *in vitro* results suggest that SIRT2 deficiency increases protein synthesis, whereas SIRT2 overexpression suppresses protein synthesis. SIRT2-deficient mice exhibit age-associated and neurohormone-induced cardiac hypertrophy. Here, we report increased global protein synthesis in the hearts of young SIRT2-deficient mice, which may contribute to the development of cardiac hypertrophy. Conversely, cardiac-specific overexpression of SIRT2 reduces global protein synthesis in mice hearts. Mechanistically, SIRT2 binds to and deacetylates Rheb at K151 residue to enhance ubiquitin-proteosome-mediated degradation of Rheb. Depletion of Rheb rescues the increased protein synthesis in SIRT2-inhibited conditions. Our findings suggest that SIRT2 activation can be a potential therapeutic for treating diseases associated with increased protein synthesis.

## Introduction

Protein synthesis and degradation are tightly regulated processes that are controlled by many regulators to maintain cellular proteostasis (Klaips, Jayaraj et al., 2018). Protein synthesis is a highly energy-consuming process that competes with other cellular processes in terms of energy consumption (Princiotta, Finzi et al., 2003). Dysregulation in protein synthesis is associated with diseases like cancer (Jia, Yuan et al., 2023, Rubio, Garland et al., 2022), neurodegenerative diseases (Chartier-Harlin, Dachsel et al., 2011, Wiebe, Nagpal et al., 2020), and cardiac diseases (Liao, Gao et al., 2021, Wang, Liao et al., 2021). Protein synthesis is one of the key cellular processes regulated by aging-related pathways (Tavernarakis, 2008), and studies suggest that down-regulation of protein synthesis can enhance the lifespan of model organisms (Syntichaki, Troulinaki et al., 2007). Reduction in global protein synthesis downregulates the accumulation of damaged and misfolded proteins, therefore improving cellular health (Kikis, Gidalevitz et al., 2010). Protein synthesis activation largely correlates with nutrient availability and cellular energy state (Jewell & Guan, 2013). NAD^+^ levels render an assessment of the cellular energetic states, where high NAD^+^ levels usually represent decreased cellular energy (Katsyuba, Romani et al., 2020). NAD^+^ and NAD^+^-dependent enzymes regulate multiple cellular processes and metabolic signaling pathways (Ying, 2006). Low levels of NAD+ are associated with the progression of various diseases, including cancers, metabolic disorders, cardiovascular disorders, and neurodegenerative diseases (Elhassan, Philp et al., 2017, Zapata-Pérez, Wanders et al., 2021). Sirtuins, a group of NAD^+^-dependent deacetylase enzymes, act as cellular energy sensors by sensing high NAD+ levels and utilize NAD^+^ to deacylate/deacetylate numerous proteins and regulate their functions (Massudi, Grant et al., 2012, Ottens, Franz et al., 2021). Seven isoforms of sirtuins (SIRT1-7) are reported in mammals; each has distinct subcellular localization and function (North & Verdin, 2004). Sirtuins regulate multiple cellular processes in cells, such as cellular metabolism, transcription, DNA repair, and protein synthesis (Michan & Sinclair, 2007). Previous findings place emphasis on the involvement of, SIRT1 (Shan, Fan et al., 2017), SIRT6 (Ravi, Jain et al., 2019), and SIRT7 (Tsai, Greco et al., 2014) in regulating global protein synthesis. However, the role of other sirtuins in this area remains unclear. SIRT2, a major cytoplasm localized sirtuin, is involved in the regulation of various cellular events like cytoskeleton organization, cellular metabolic activities, inflammation, and nuclear membrane dynamics (de Oliveira, Sarkander et al., 2012). However, being the only cytoplasmic sirtuin, where most of the translational machinery is present, how SIRT2 contributes to regulating protein synthesis remains unexplored.

The mechanistic Target of Rapamycin (mTOR) is one of the master controllers of protein synthesis in cells (Saxton & Sabatini, 2017b). mTOR is a protein kinase that belongs to the PI3K-related kinase family and the main constituent of two distinct protein complexes known as mTORC1 and mTORC2; these complexes have different functions, with mTORC1 governing global protein synthesis (Saxton & Sabatini, 2017b, Wullschleger, Loewith et al., 2006). Ras homolog enriched in the brain (Rheb), a small GTPase of the Ras Superfamily, is crucial for mTORC1 activation (Long, Lin et al., 2005), In the absence of growth stimulation, Rheb remains bound with the TSC1/2 complex. TSC2 acts as Rheb GAP and induces intrinsic GTPase activity of Rheb to keep it inactive in Rheb-GDP form (Zhang, Gao et al., 2003). Upon receiving adequate growth stimulatory signals, Akt-dependent multisite phosphorylation of TSC2 releases the TSC1/2 complex from Rheb leading to the Rheb activation. Active Rheb in its GTP-bound form recruits mTORC1 on the lysosomal membrane and activates mTOR kinase activity within the mTORC1 complex (Garami, Zwartkruis et al., 2003, Long et al., 2005, Saxton & Sabatini, 2017a). Active mTORC1, with the help of its accessory subunit raptor, phosphorylates distinct mTORC1 downstream targets like S6 kinase and 4EBP1. This further results in increased protein synthesis (Morita, Gravel et al., 2015, Saxton & Sabatini, 2017a), majorly due to increased translation of cap-dependent mRNAs (Dibble & Manning, 2013, Le Sage, Cinti et al., 2016, Ma & Blenis, 2009).

Previous reports suggest that protein synthesis is upregulated during cardiac hypertrophy to meet the immense protein demand of the hypertrophic heart (Grund, Szaroszyk et al., 2019, Yan, Tang et al., 2021). The hypertrophic heart often exhibits enhanced activation of mTORC1 signaling, and inhibition of the mTORC1 pathway improves cardiac function in mice models via reducing cardiac hypertrophy and fibrosis (Gao, Wong et al., 2006). Current knowledge about sirtuins provides insight into their possible roles in cardiac health (Ravi, Mishra et al., 2021). Our recent study on SIRT6 has elucidated the importance of SIRT6 in the regulation of protein synthesis in the context of cardiac hypertrophy (Ravi et al., 2019). Recent findings show that SIRT2 levels decrease in the cardiac tissues of aged mice, and *Sirt2*-KO mice develop aging-associated hypertrophy (Tang, Chen et al., 2017). Moreover, *Sirt2*-KO mice display increased sensitivity towards neurohormonal stress-induced cardiac hypertrophy and fibrosis (Sarikhani, Maity et al., 2018a, Tang et al., 2017). These recent reports emphasize the importance of understanding if SIRT2 plays role in regulating global protein synthesis.

In the current study, we find that SIRT2 plays a key role in regulating global protein synthesis inside cells by deacetylating Rheb and facilitating its proteosome-mediated degradation. In our current study, we have used the *Sirt2*-KO mice model to understand the significance of increased Rheb levels, mTORC1 activity, and upregulated global protein synthesis in the development of cardiac hypertrophy in *Sirt2*-KO mice heart.

## Results

### SIRT2 negatively regulates global protein synthesis

Our previous findings showed that SIRT2 levels are downregulated in the cardiac tissue of mice injected with cardiac stress-inducing drugs Isoproterenol (ISO) and Phenylephrine (PE) (Sarikhani et al., 2018a). The rate of protein synthesis is also reported to be upregulated in the heart of mice that develop ISO-induced hypertrophy (Sarikhani et al., 2018a). Therefore, we hypothesized that SIRT2 might have a role in regulating global protein synthesis. To determine if SIRT2 directly regulates protein synthesis, we inhibited SIRT2 using AGK2 (SIRT2 inhibitor). We observed that protein synthesis rates were significantly upregulated upon SIRT2 inhibition, as confirmed by both Western and confocal-based assessments of puromycin incorporation under SIRT2 inhibition, suggesting a direct involvement of SIRT2 **[Figure 1A-E]** in regulating protein translation. We observed a significantly enhanced rate of protein synthesis in cells that were transiently depleted for SIRT2 using SIRT2-specific siRNA **[Figure 1F, H]**. Since we observed that inhibiting SIRT2 activity or depleting its endogenous levels could significantly enhance the rate of protein synthesis, we further confirmed if SIRT2 overexpression could reduce protein synthesis. Transiently overexpression of SIRT2 led to a significantly lower rate of protein synthesis **[Figure 1G, I].** SIRT2 is known to remove acetylation (Kim, Vassilopoulos et al., 2011) or long-chain acylation (Jing, Zhang et al., 2017) from the lysine residues of its target proteins. We transiently overexpressed SIRT2 catalytically inactive mutant (SIRT2 N168A) to check if SIRT2-mediated regulation of protein synthesis depends on its catalytic activity. Interestingly, we observed that the rate of protein synthesis was significantly higher in cells overexpressing SIRT2 N168A mutant than in cells expressing the pcDNA control **[Figure 1J, K]**. This observation suggests a possible dominant negative effect of SIRT2 N168A mutant as SIRT2 dimerization is crucial for SIRT2 deacetylase activity (Yang, Nicely et al., 2023); previous studies have also reported SIRT2 N168A mutant as catalytically inactive dominant mutant (Li, Xu et al., 2008). We also checked if and how SIRT2 responds during complete cellular starvation, a state where protein synthesis remains inhibited, we treated cells with EBSS (Earle’s Balanced Salt Solution) and observed that complete cell starvation increases SIRT2 levels and activity concomitant with decreased global protein synthesis **[Figure S1A, B]**. These findings suggest that SIRT2 is a negative regulator of protein synthesis.

**Figure 1.**
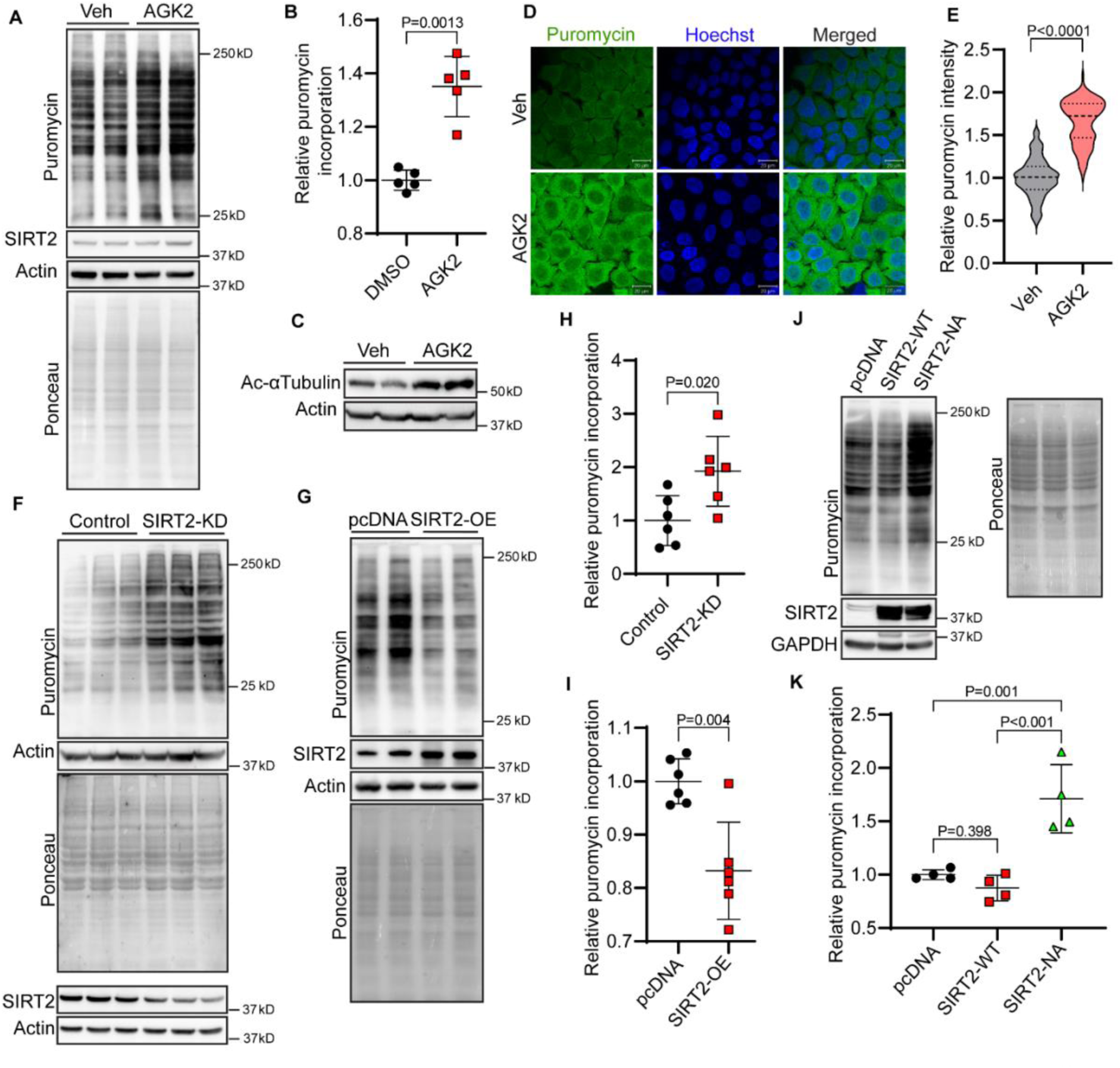
SIRT2 negatively regulates global protein synthesis. **A.** Representative western blot images of SUnSET analysis depicting changes in protein synthesis rate in HeLa cells treated with DMSO or SIRT2 inhibitor AGK2. **B.** Quantification of puromycin incorporation depicted in Figure 1A. The results are expressed as the fold change relative to DMSO-treated controls, n = 5. P values shown are from unpaired, 2-tailed Student’s t-test. Data are presented as mean ± s.d, **C.** Representative western blot images showing changes in acetyl tubulin levels in Hela cells after DMSO (Vehicle) and AGK2 treatment. **D.** Representative images of IFC—SUnSET analysis in HeLa cells treated with vehicle (DMSO) and AGK2. Puromycin staining is shown in green, Nuclei are stained with Hoechst 33342 and shown in blue. **E.** Quantitative representation of IFC-SUnSET analysis shown in figure 1D, Puromycin intensity was normalized against vehicle, n=90-100 cells per group. P values shown are from the Mann-Whitney test. data are shown as median with X and Y percentile. **F.** Representative images of Western blotting SUnSET analysis depicting the changes in protein synthesis rate in control and SIRT2 depleted (SIRT2-KD) HeLa cells. SIRT2 was depleted using SIRT2-specific siRNA, and scrambled siRNA was used as a control. Separate SDS-PAGE was run, and SIRT2 knockdown was confirmed via immunoblotting **G.** Representative images of Western blotting SUnSET analysis depicting the changes in protein synthesis rate in control and SIRT2 overexpressing HeLa cells. HeLa cells were transfected with either pcDNA (control) or SIRT2-WT plasmids, and SIRT2 overexpression was confirmed via immunoblotting. **H.** Quantitative representation of puromycin incorporation depicted in Figure 1F. The values are shown as the fold change relative to scrambled siRNA-treated control groups. n = 6. P values shown are from unpaired, 2-tailed Student’s t-test. Data are presented as mean ± s.d, **I.** Quantitative representation of puromycin incorporation depicted in Figure 1G. The results are expressed as the fold change relative to pcDNA transfected control groups. n = 6. P values shown are from unpaired, 2-tailed Student’s t-test. Data are presented as mean ± s.d, **J.** Representative images of western blotting SUnSET analysis in HeLa cells transfected with pcDNA, SIRT2-WT, or SIRT2-N168A plasmids. The plasmids were transfected for 48 h, and the expression of SIRT2 and its catalytic mutant were confirmed by Western blotting. **K.** Quantitative representation of puromycin incorporation depicted in Figure 1L. The results are expressed as the fold change relative to pcDNA transfected control groups. n = 4. P values shown are from ordinary one-way ANOVA with Holm-Sidak’s multiple comparisons test. Data are presented as mean ± s.d,

These findings suggest that SIRT2 is a negative regulator of protein synthesis.

### SIRT2 affects protein synthesis by regulating mTOR kinase activity

Most mRNAs in eukaryotic cells undergo translation through a process known as CAP-dependent translation (Sonenberg & Hinnebusch, 2009). Since cap-dependent translation is the primary translatory pathway in cells, we were interested in understanding if SIRT2 regulates global protein synthesis via the regulation of the cap-dependent translation process. To test this, we used the EMCV-BICIS construct, a dual luciferase reporter vector that can be used to measure cap-dependent and cap-independent translation separately. The Renilla/firefly ratio provides a ratio of cap-dependent translation in cells (Choo, Yoon et al., 2008, Dave, George et al., 2019) **[Figure 2A]**. Significant upregulation was observed in cap-dependent translation in cells upon SIRT2 inhibition using AGK2 **[Figure 2B]**. AGK2-mediated SIRT2 inhibition was confirmed by increased acetylation of α-tubulin **[Figure 2E]**. Furthermore, we also observed a significant increase in cell cap-dependent translation upon SIRT2 knockdown **[Figure 2C]**. We found a reduction in cap-dependent translation under SIRT2 overexpression, whereas overexpression of catalytically inactive mutant SIRT2 N168A did not show such a reduction in cap-dependent translation **[Figure 2D]**. Initiation of translation in a cap-dependent manner is the most evaluated regulatory step that depends on recognizing the 5’ cap region by the eIF4F complex. This complex consists of cap-binding factor eIF4E, scaffold protein eIF4G, and a helicase eIF4A (Gandin, English et al., 2022). Previous reports have shown that the assembly of the eIF4F complex is largely governed by mTORC1(Gingras, Gygi et al., 1999, Sonenberg & Hinnebusch, 2009). mTORC1 activation often correlates with the sufficiency of dietary intake, and the activation is inhibited during fasting or limited energy source availability. Activated mTOR kinase phosphorylates downstream targets like S6 kinase and 4EBP1, which leads to an upregulation of protein translation, increasing mostly the translation of cap-dependent mRNAs (Le Sage et al., 2016). Previous studies have shown that phosphorylation of Ser-2448 at mTOR is dependent on mTOR-kinase activity (Bonnet, Palandri et al., 2024, Chiang & Abraham, 2005, Ochiai, Shima et al., 2024, Ravi et al., 2019). Anabolic factors such as insulin, nutrients, and amino acids can upregulate Ser-2448 phosphorylation (Bonnet et al., 2024, Chiang & Abraham, 2005). To understand whether SIRT2 regulates protein synthesis through the regulation of the mTOR pathway, we treated SIRT2-depleted cells with mTOR inhibitors, Torin1 and Rapamycin, and observed that cells treated with these inhibitors do not exhibit an increase in protein synthesis rate **[Figure 2F]**. Upon SIRT2 inhibition, concomitant with the previously observed increased cap-dependent translation, phosphorylation of mTOR was significantly upregulated **[Figure 2G, H]**. Increased p-mTOR levels were further observed under the SIRT2 knocked-down condition **[Figure 2I, J]**. These findings strongly indicate that SIRT2 regulates protein synthesis by regulating mTOR kinase activity.

**Figure 2.**
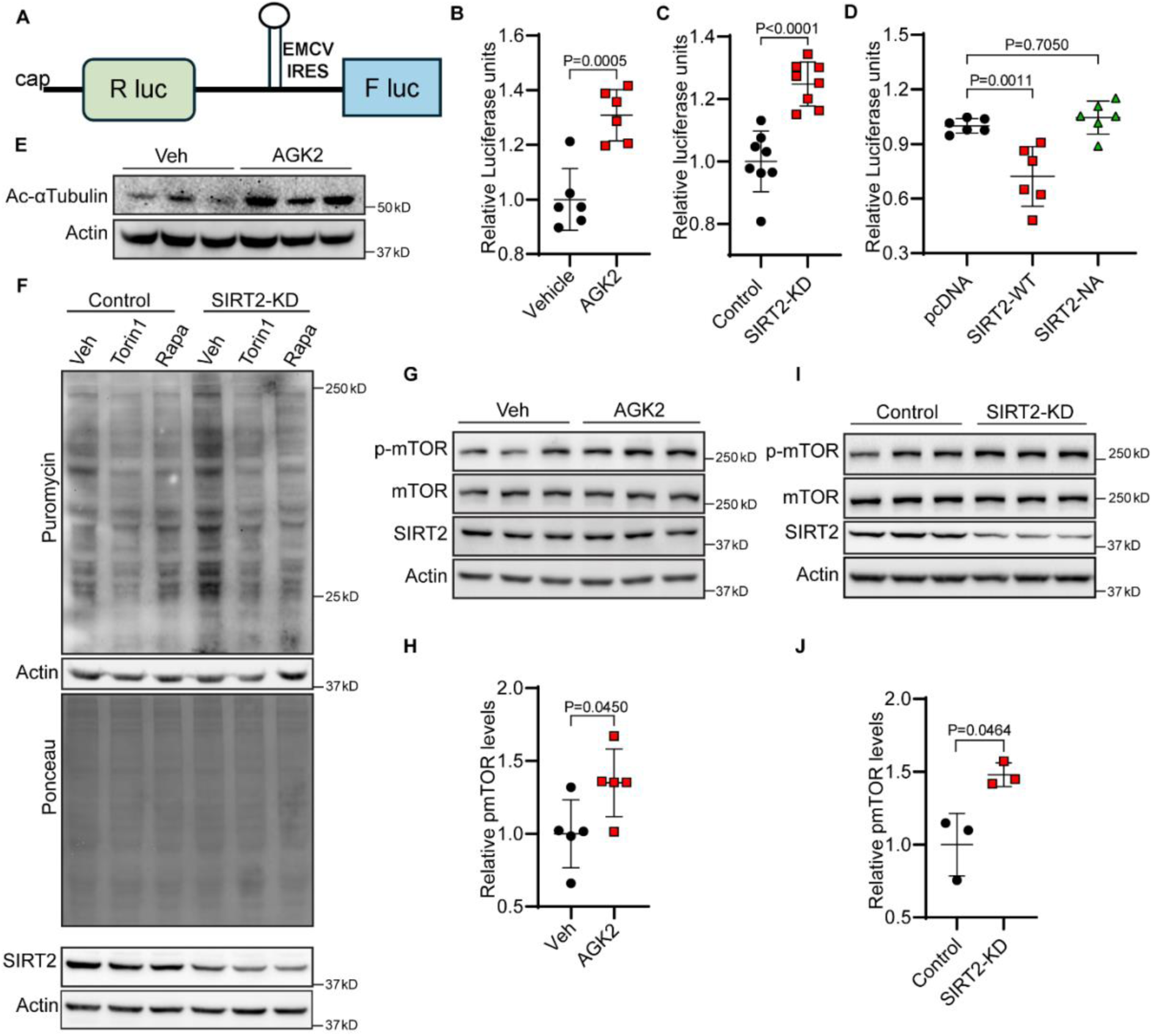
SIRT2 affects protein synthesis by regulating mTOR kinase activity. **(A)** Schematic representation of luciferase reporter EMCV-BICIS construct for assessing cap-dependent translation. **(B)** Quantitative representation of Cap-dependent translation measured using luciferase reporter EMCV-BICIS construct in HeLa cells treated with either DMSO (Vehicle) or AGK2. The results are expressed as the fold change relative to control group. P values shown are from unpaired, 2-tailed Student’s t-test. Data are presented as mean ± s.d, n=6. **(C)** Quantitative representation of Cap-dependent translation measured using luciferase reporter EMCV-BICIS construct in HeLa cells transfected with either Control or SIRT2 specific siRNA. The results are expressed as the fold change relative to control group. P values shown are from unpaired, 2-tailed Student’s t-test. Data are presented as mean ± s.d, n=8. **(D)** Quantitative representation of Cap-dependent translation rates measured using luciferase reporter EMCV-BICIS construct in control, SIRT2-WT, and SIRT2 N168A mutant overexpressing HeLa cells. The results are expressed as the fold change relative to control group. P values shown are from ordinary one-way ANOVA with Dunnett’s multiple comparisons test. Data are presented as mean ± s.d, n=6. **(E)** Representative images of western blotting analysis showing changes in Acetyl tubulin levels in Hela cells after DMSO (Vehicle) and AGK2 treatment. **(F)** Representative images of Western blotting SUnSET analysis depicting the changes in protein synthesis rate in control and SIRT2 depleted (SIRT2-KD) HeLa cells treated with either vehicle or mTOR inhibitors Torin1 and Rapamycin. SIRT2 was depleted using SIRT2-specific siRNA, and Scrambled siRNA was used as a control. Separate SDS-PAGE was run, and SIRT2 knockdown was confirmed via immunoblotting. n=3. **(G)** Representative western blotting images depicting changes in mTOR activation (mTOR phosphorylation) in HeLa cells treated with DMSO (Vehicle) or SIRT2 inhibitor AGK2. **(H)** Quantitative representation of mTOR activation depicted in Figure 2G. The results are expressed as the fold change relative to vehicle-treated controls. P values shown are from unpaired, 2-tailed Student’s t-test. Data are presented as mean ± s.d, n=5. **(I)** Representative western blotting images depicting changes in mTOR activation (mTOR phosphorylation) in control and SIRT2-depleted HeLa cells. **(J)** Quantitative representation of mTOR activation depicted in Figure 2I. The results are expressed as the fold change relative to Scrambled siRNA transfected controls. P values shown are from unpaired, 2-tailed Student’s t-test. Data are presented as mean ± s.d, n=3.

### SIRT2 deacetylates Rheb GTPase and promotes its proteasomal degradation

SIRT2 primarily resides in the cytoplasm and regulates crucial cellular processes by post-translationally modifying its various substrate proteins. Since SIRT2 affects protein translation and mTOR activity, we were interested in identifying novel substrates of SIRT2 that belong to the mTOR signaling pathway. Recent findings suggest that SIRT2 can deacylase multiple small GTPases inside cells to protect against Shigella infection, and Rheb GTPase is one of the substrates for SIRT2 deacylation during Shigella infection, where effector molecules from *Shigella* reportedly fatty acylates lysine residues of Rheb and other small GTPases (Wang, Zhang et al., 2022). A recent study on SIRT2 interactome in HeLa cells also suggests a possible interaction of SIRT2 with the Rheb protein (Eldridge, Pereira et al., 2020). Therefore, we verified if SIRT2 could directly interact with Rheb and deacylate Rheb. Co-immunoprecipitation showed that endogenous Rheb interacts with SIRT2 even at basal conditions **[Figure 3A]**. Since SIRT2 was previously reported for removing Rheb’s lysine fatty acylation during *Shigella* infection, we were interested to study if SIRT2 could also mediate its regulation of mTOR signalling by removing the fatty acylation on Rheb protein. To do so, we first used a click chemistry-based approach. Flag-tagged Rheb was overexpressed in control and SIRT2 knockdown (KD) HEK 293T cells. Alk14 was used to label acylated proteins in cells, and acylation level could be observed using in-gel fluorescence after click chemistry to conjugate a rhodamine fluorescent dye. Using this method, we found that Rheb was not acylated in either the control or the SIRT2 KD cells. This suggests that in a normal healthy cellular environment when there is no *Shigella* infection, the Rheb protein is not acylated **[Figure S2A]**. Previous reports suggest a possible role of Rheb acetylation in mTORC1 activation in multiple cells, where acetylated FKBP12 interacts with acetylated Rheb and facilitates mTORC1 kinase activity (Hu, Chen et al., 2021). However, it remains unknown how Rheb acetylation is regulated and how the acetylation of Rheb promotes its ability to induce mTORC1 activation. Therefore, we hypothesize that SIRT2 may regulate Rheb-mediated mTORC1 activation by deacetylating Rheb.

**Figure 3.**
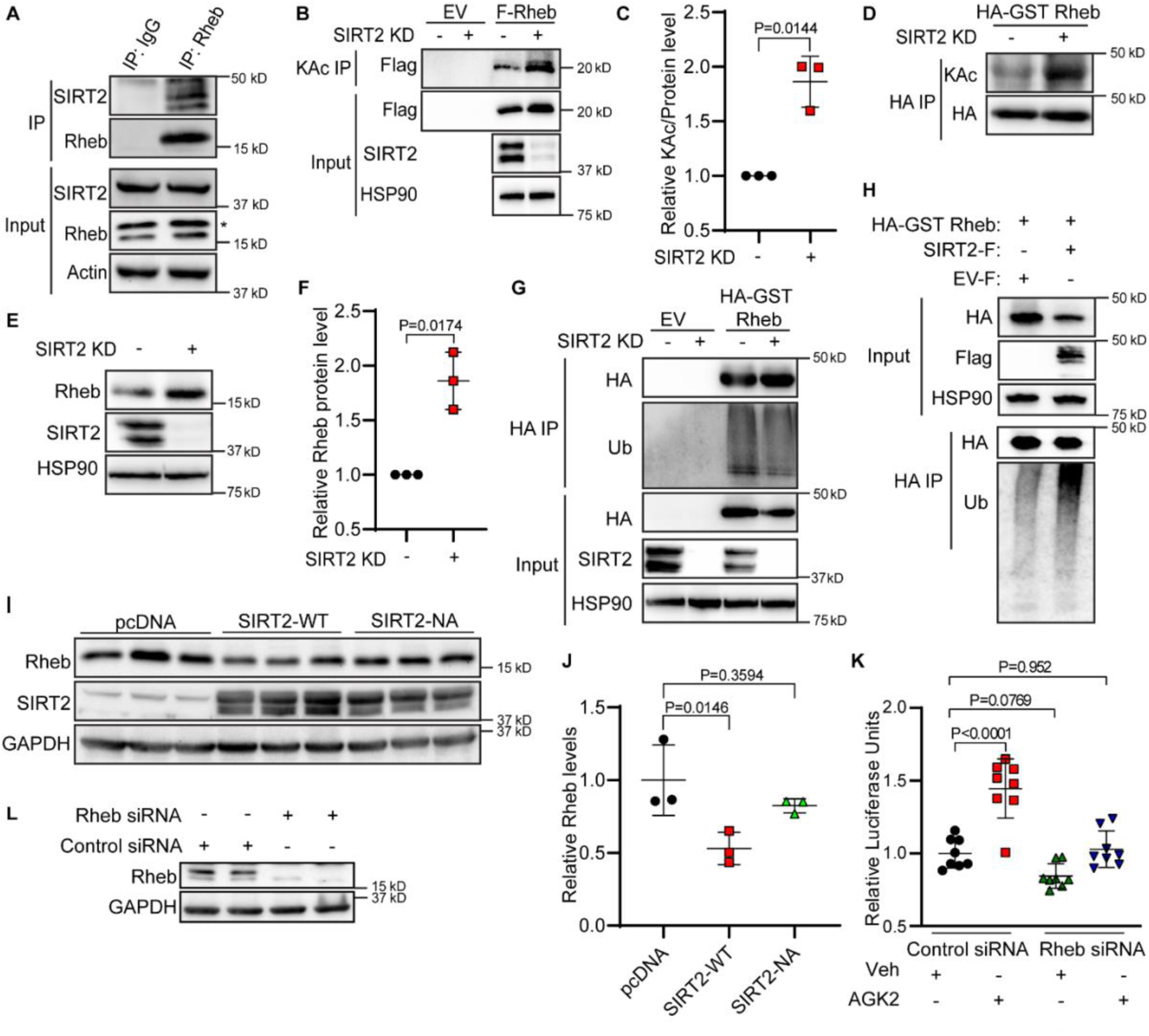
SIRT2 deacetylates Rheb GTPase and promotes its proteasomal degradation. **(A)** Representative immunoblot images showing the interaction of SIRT2 with Rheb in HeLa cells. Rheb was immunoprecipitated and probed for SIRT2. Whole cell lysate (WCL) was also immunoblotted to check the loading. n = 3 **(B)** Rheb is deacetylated by SIRT2. Flag-tagged Rheb or Flag-tagged empty vector was transfected into control and SIRT2 knockdown HEK 293T cells. Acetylation was determined using acetyl lysine pull-down and western blot for Flag-Rheb. **(C)** Quantification of Rheb acetylation (normalized to protein level). P values shown are from ratio paired Student’s t-test. Data are presented as mean ± s.d, n=3. **(D)** Rheb is deacetylated by SIRT2. HA-GST Rheb was transfected into control and SIRT2 knockdown HEK 293T cells. After HA IP pull down, lysine acetyl antibody was used to detect acetylated Rheb in western blot. **(E)** Endogenous Rheb protein level increases when SIRT2 is knocked down. Endogenous Rheb levels in control and SIRT2 knockdown HEK 293T cells were evaluated using western blot. **(F)** Relative Rheb to total protein level ratios were quantified. P values shown are from ratio paired, 2-tailed Student’s t-test. Data are presented as mean ± s.d, n=3. **(G)** Knockdown of SIRT2 decreases Rheb ubiquitination. HA-GST Rheb was expressed in control and SIRT2 knockdown HEK 293T cells and was pulled down by HA beads. The ubiquitination level was analyzed using an anti-ubiquitin western blot. **(H)** Overexpression of SIRT2 increases Rheb ubiquitination. Flag-tagged SIRT2 or empty vector, along with HA-GST Rheb, were transfected into HEK 293T cells. Rheb was pulled down by HA beads, and ubiquitination levels were analyzed using Western blot. **(I)** Representative western blot images depicting changes in Rheb protein levels in control Hela cells or cells overexpressing SIRT2-WT or SIRT2 N168A mutant. **(J)** Quantification of Rheb protein levels depicted in Figure 3I. The results are expressed as the fold change relative to control group. P values shown are from the Kruskal-Wallis test for nonparametric data with Dunn’s multiple comparisons test. Data are presented as mean ± s.d, n = 3. **(K)** Quantification of Cap-dependent translation measured using luciferase reporter EMCV-BICIS construct in HeLa cells transfected with scrambled siRNA or Rheb siRNA and treated with either vehicle (DMSO) or AGK2. The results are expressed as the fold change relative to control group. P values shown are from ordinary one-way ANOVA with Dunnett’s multiple comparisons test. Data are presented as mean ± s.d, n=8. **(L)** Representative western blot images depicting changes in Rheb protein levels in HeLa cells transfected with scrambled siRNA or Rheb siRNA.

To check whether Rheb is a deacetylation substrate of SIRT2, we overexpressed Flag-tagged Rheb in control and SIRT2 KD HEK 293T cells. Anti-acetyl lysine beads were used to pull down the acetylated proteins in the sample, and an anti-Flag western blot was used to analyze the acetylation level of Rheb. The acetylation level of Rheb increased in SIRT2 KD cells, indicating that SIRT2 could deacetylate Rheb (**Figure 3B-C**). We also pulled down Rheb and detected its acetylation level by anti-acetyl lysine western blot. Again, Rheb was more acetylated in SIRT2 KD cells than in control cells (**Figure 3D**). Thus, our data support the idea that Rheb is a deacetylation substrate of SIRT2. Interestingly, total protein levels of Rheb were also significantly upregulated in SIRT2 KD cells **[Figure 3E, F]**. This observation suggests a possible role of SIRT2 in regulating Rheb’s stability. Multiple studies have shown that lysine acetylation can promote protein stability and decrease protein ubiquitination (Buchwald, Krämer et al., 2009, Inuzuka, Gao et al., 2012, Nihira, Ogura et al., 2017, Shimizu, Gi et al., 2021). To check if lysine acetylation of Rheb facilitates Rheb stability, we examined Rheb ubiquitination in control and SIRT2 KD cells. In line with increased Rheb protein levels, we observed that SIRT2 KD cells exhibit decreased ubiquitination of Rheb compared to control KD cells **[Figure 3G]**. Furthermore, Rheb ubiquitination increased upon SIRT2 overexpression **[Figure 3H]**. Overexpression of wild-type SIRT2 led to decreased Rheb protein levels, whereas overexpression of SIRT2 N168A mutant did not **[Figure 3I, J]**. Furthermore, we observed that downregulating Rheb can rescue the increased cap-dependent translation under SIRT2-inhibited conditions. **[Figure 3K, L].** These findings indicate that SIRT2 deacetylates Rheb and promotes its ubiquitination and degradation.

### Lysine 151 residue in the Rheb is the site for SIRT2-mediated deacetylation, and acetylation of lysine 151 stabilizes the Rheb

To mark the lysin residues in Rheb, targeted by SIRT2 for deacetylation, we performed a mass-spectrometric analysis in cells treated with AGK2 (SIRT2 inhibitor). Our mass spectrometry data reveals lysine 151 (K151) of Rheb protein as a possible acetylated residue under SIRT2 inhibition, suggesting that SIRT2 may deacetylate Rheb at the K151 position **[Figure 4 A]**. Previous reports suggest that SIRT2 regulates the K147 acetylation status of a small GTPase, KRas (Song, Biancucci et al., 2016). Interestingly, Rheb and KRas share a similar sequence stretch of amino acids for K151 and K147 residues, respectively **[Figure S3 A]**. Moreover, our sequence alignment data suggest that lysine K151 residue is conserved across different species **[Figure S3 B]**. Structural analysis of Rheb Shows that K151 is located near the GTP binding pocket of Rheb **[Figure 4B]**. To understand the structural and functional significance of K151 acetylation, we performed site-directed mutagenesis in pRK5 HA-GST Rheb construct to generate Rheb K151R (mimicking deacetylated form) and Rheb K151Q (mimicking acetylated form) mutants of Rheb. HA-GST Rheb (Rheb WT) and Rheb K151R were expressed in SIRT2 KD cells and were pulled down using anti-HA beads. Acetylation blot showed a consistent and significant decrease in acetylation levels of Rheb K151R compared to Rheb WT, confirming that SIRT2 deacetylates Rheb at K151 **[Figure 4C, D]**. Moreover, when the western blot of the same samples was blotted with an anti-ubiquitin antibody, increased ubiquitination was observed in the K151R mutant **[Figure 4C]**. To further confirm that it was indeed the deacetylation that was responsible for the increase in Rheb ubiquitination, we overexpressed Rheb WT, K151R mutant, or K151Q mutant in control and SIRT2 knockdown cells, similar to previous observations, The K151R mutant exhibited significantly increased Rheb ubiquitination, while the K151Q mutant, which mimics the acetylated form of Rheb, did not increase Rheb ubiquitination **[Figure E]**. Our data confirmed that SIRT2 deacetylates Rheb at K151 and promotes Rheb ubiquitination and degradation.

**Figure 4.**
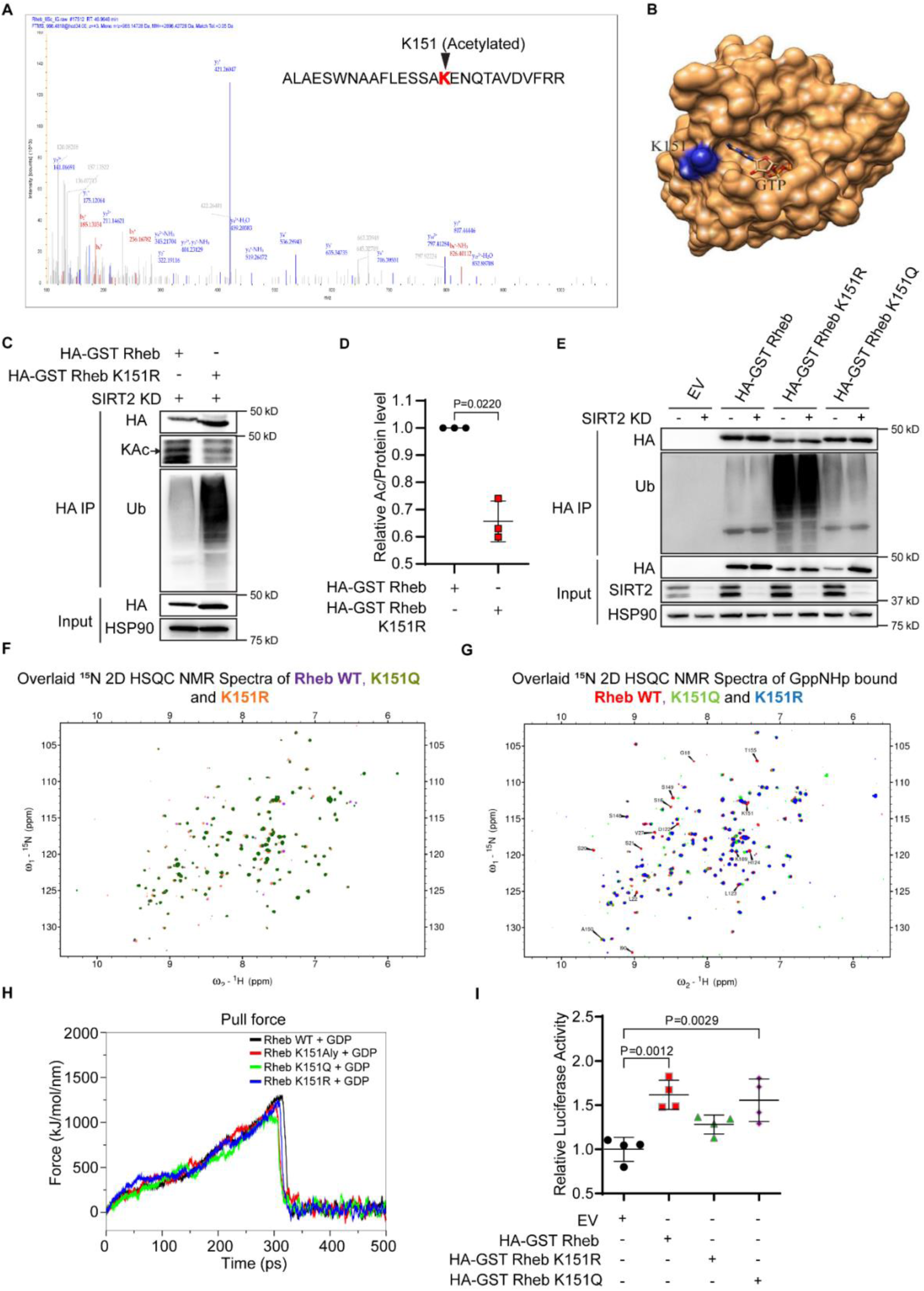
Lysine 151 residue in the Rheb is the site for SIRT2-mediated deacetylation, and acetylation of lysine 151 stabilizes the Rheb. **(A)** LC/MS spectra depicting acetylation of lysine residue 151 (K151) in Rheb protein under AGK2 treatment in HeLa cells. **(B)** 3D structure of GTP bound Rheb depicting the position of K151 residue **(C)** SIRT2 deacetylates Rheb at K151, which in turn increases Rheb’s ubiquitination. HA-GST Rheb or HA-GST Rheb K151R were transfected into SIRT2 knockdown HEK 293T cells and pulled down with HA IP beads. Acetylation and ubiquitination were detected using western blots. **(D)** Relative Rheb acetylation levels (normalized to Rheb protein levels) were quantified and compared between HA-GST Rheb and HA-GST Rheb K151R in SIRT2 knockdown HEK 293T cells. P values shown are from ratio paired, 2-tailed Student’s t-test. Data are presented as mean ± s.d, n = 3. **(E)** Deacetylation of Rheb at K151 by SIRT2 increases ubiquitination. HA-GST Rheb, HA-GST Rheb K151R, and HA-GST Rheb K151Q were transfected into control or SIRT2 knockdown HEK 293T cells. The ubiquitination level of Rheb was analyzed by HA IP, and western blot. **(F)** Representative overlaid 2D ^1^H-^15^N HSQC NMR spectra of Rheb WT, K151Q, and K151R Rheb mutants. **(G)** Representative overlaid 2D ^1^H-^15^N HSQC NMR spectra of GppNHp bound Rheb WT, K151Q, and K151R Rheb mutants. **(H)** Graphical depiction of Pulling force simulation analysis showing the strength of GDP association with Rheb WT, Rheb K151Aly, and K151R, K151Q Rheb mutants. **(I)** Quantitative representation of Cap-dependent translation measured using luciferase reporter EMCV-BICIS construct in HeLa cells overexpressing HA-GST Rheb, HA-GST Rheb K151R, and HA-GST Rheb K151Q mutants. The results are expressed as the fold change relative to the Empty vector (EV) transfected control group. P values shown are from ordinary one-way ANOVA with Tukey’s multiple comparisons test. Data are presented as mean ± s.d, n=4.

We used NMR spectroscopy to characterize the structural significance of these mutants. 2D ^1^H-^15^N HSQC NMR spectra were recorded for Rheb WT, Rheb K151Q, and Rheb K151R in basal and non-hydrolyzable GTP analog, GppNHp bound form. Our analysis showed that K151R or K151Q mutation does not affect the overall structure or the GTP binding ability of Rheb **[Figure 4F, G, S4 A-D]**. GDP dissociation inhibitors are a class of factors that inhibit GDP-GTP exchange and regulate the activity of multiple small GTPases (Collins, 2003, DerMardirossian & Bokoch, 2005, Han, Jeong et al., 2021); to understand if acetylation of Rheb at K151 has any role in GDP release from Rheb, we performed Pulling force simulations to investigate the GDP binding strength in Rheb WT, Rheb K151acetylated, and mutants (K151R, K151Q). We did not observe any significant difference in the pulling force required to remove GDP among these groups **[Figure 4H**, **Table 1],** suggesting that acetylation of Rheb at K151 might not affect GTP binding and GDP dissociation. Next, we assessed the ability of these mutants to induce cap-dependent translation; we observed a significant increase in cap-dependent translation in cells transfected with Rheb WT, and K151Q mutant when compared with the control group, whereas the overexpression of the K151R mutant did not induce cap-dependent translation **[Figure 4I, S4E]**. These findings suggest that SIRT2 deacetylates Rheb by targeting K151 residue and facilitates Rheb ubiquitination and degradation, inhibiting Rheb-mediated mTORC1 activation.

### Acetylation of Rheb at K151 does not affect its membrane binding

We have performed all-atom molecular dynamics simulations to understand the effect of c-terminal farnesylation for wildtype as well as mutations that serve as acetylated or deacetylated mimics on membrane association. To gain molecular insights, protein-bilayer systems have been simulated. Four different systems, i.e. (a) WT Rheb with bilayer, (b) K151Acetylated with bilayer, (c) K151Q as an acetylated mimic with bilayer, and (d) K151R as a deacetylated mimic with bilayer have been set up. To cancel any bias generated due to starting orientation, all four systems have been started with the same initial orientation of the protein on the bilayer **[Figure 5A]**. For mimicking the membrane in simulation, 320 POPC and 96 POPS in ratio of 80:20 have been used as bilayer composition. Simulation outputs were analyzed by a simple Z-distance calculation between the Cα atoms of each residue with the z-coordinate of the phosphate plane of the bilayer. If the z-distance is less than zero, it means membrane association and vice versa. The analysis has been performed over 200 snapshots of last 20 ns run. Interestingly, there is no significant effect of mutation/acetylation mimic on membrane association **[Figure 5B]**. Though the traces of membrane association have been observed in the K151Q-bilayer simulations. Moreover, to confirm the membrane association, we have calculated the z-distance between the side chain atom of the 172nd residue and the phosphate plane **[Figure 5C]**. It clearly shows that the association is quite transient. Thereby, confirming no significant impact of these acetylation upon membrane association.

**Figure 5.**
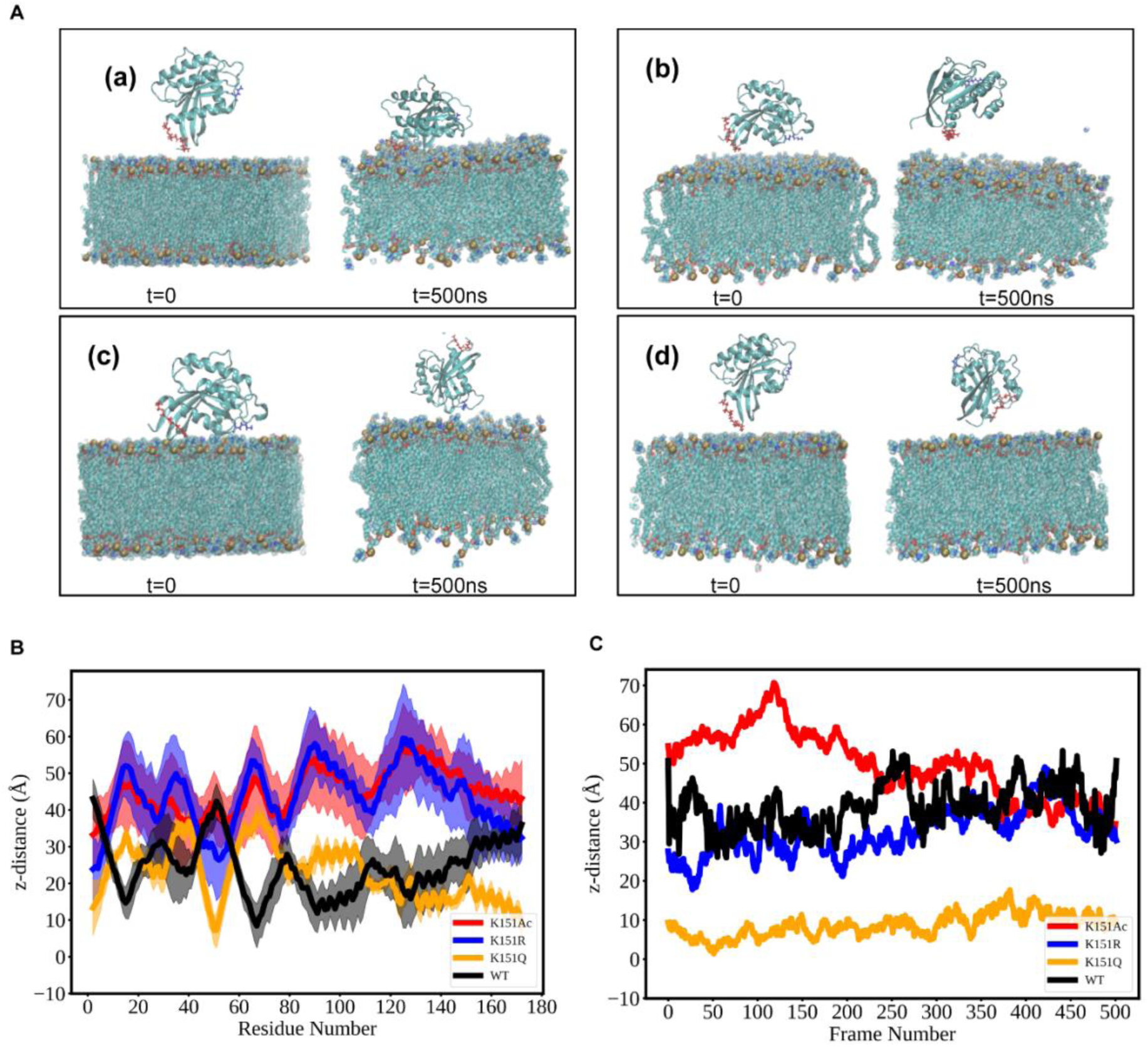
Molecular Simulation for studying the effect of acetylation upon C-terminal farnesylated Rheb protein on membrane association. **(A)** Schematic diagram representing Rheb-bilayer orientations in different protein states considered for the study. initial at t=0 and final at t=500ns. (a) WT Rheb with bilayer, (b) K151Acetylated with bilayer, (c) K151Q as acetylated mimic with bilayer, and (d) K151R as deacetylated mimic, respectively. Rheb protein has been shown in a cartoon representation, and K151 residue has been marked in bonds represented in blue color. The Farnesyl anchor is shown in red. Membrane phosphate residues has been shown in vdw representation. Water and ions have not been shown for clarity of image. **(B)** Graphical diagram representing the z-distance between the C-alpha of each residue and the phosphate plane of the bilayer. **(C)** Graphical diagram representing z-distance between 172 residue and phosphate plane of the bilayer.

### *Sirt2*-KO mice hearts exhibit hyperactivated mTOR kinase activity

Previous reports suggest that *Sirt2*-KO mice develop cardiac hypertrophy in an age-dependent manner, and *Sirt2*-KO mice are more sensitive to neurohormonal-induced cardiac hypertrophy (Sarikhani et al., 2018a, Tang et al., 2017). As mentioned earlier, increased protein synthesis is a well-known hallmark of mice hearts exhibiting cardiac hypertrophy. Cardiomyocytes develop various mechanisms to ensure the increased demand for protein in the context of increased cardiomyocytes when the heart is in the hypertrophic stage. We used this model to understand if *Sirt2*-KO mice hearts are prone to develop hypertrophy because of the upregulated protein synthesis in the absence of SIRT2. To check this, we first used young *Sirt2*-KO mice of 4 months, whose heart functions are as normal as control mice **[Figure S5 A-G]**. We tested for the protein synthesis levels in these young *Sirt2*-KO mice heart tissue using the SUnSET assay. The protein synthesis was significantly upregulated in the heart tissue of *Sirt2*-KO mice compared to WT controls **[Figure 6A, B]**. Since our *in vitro* results suggest that SIRT2 promotes Rheb degradation, we checked for Rheb levels in *Sirt2*-KO mice heart tissue. We found that Rheb protein levels were upregulated in these tissues compared to WT controls **[Figure 6C, D]**. Furthermore, mTOR signaling activation was assessed in *Sirt2*-KO mice’s heart tissues by assessing the levels of p-mTOR and p-p-70S6 Kinase 1 (p-p-70S6K1), and we found that activation of mTOR signaling was upregulated in *Sirt2*-KO mouse hearts **[Figure 6E]**. Moreover, when detecting protein synthesis levels in the heart tissues of 9-month-old *Sirt2*-KO mice, where heart tissues showed hypertrophic changes compared to wildtype controls **[Figure 6F-I, Figure S5 H-L],** protein synthesis was found to be upregulated in these mice’s heart tissues **[Figure 6J]**. Since increased protein synthesis in cardiomyocytes correlates with increased cardiomyocyte size during hypertrophic changes, we performed WGA staining in 9-month-old *Sirt2*-KO mice heart tissue sections. We found that *Sirt2*-KO mice display an increase in cardiomyocyte cross-sectional area compared to wild-type controls **[Figure 6K, L]**. These findings suggest that enhanced protein synthesis in *Sirt2*-KO mice, even at the early stage, can be a crucial factor for developing cardiac hypertrophy in *Sirt2*-KO mice when they age **[Figure S5 M-T]**.

**Figure 6.**
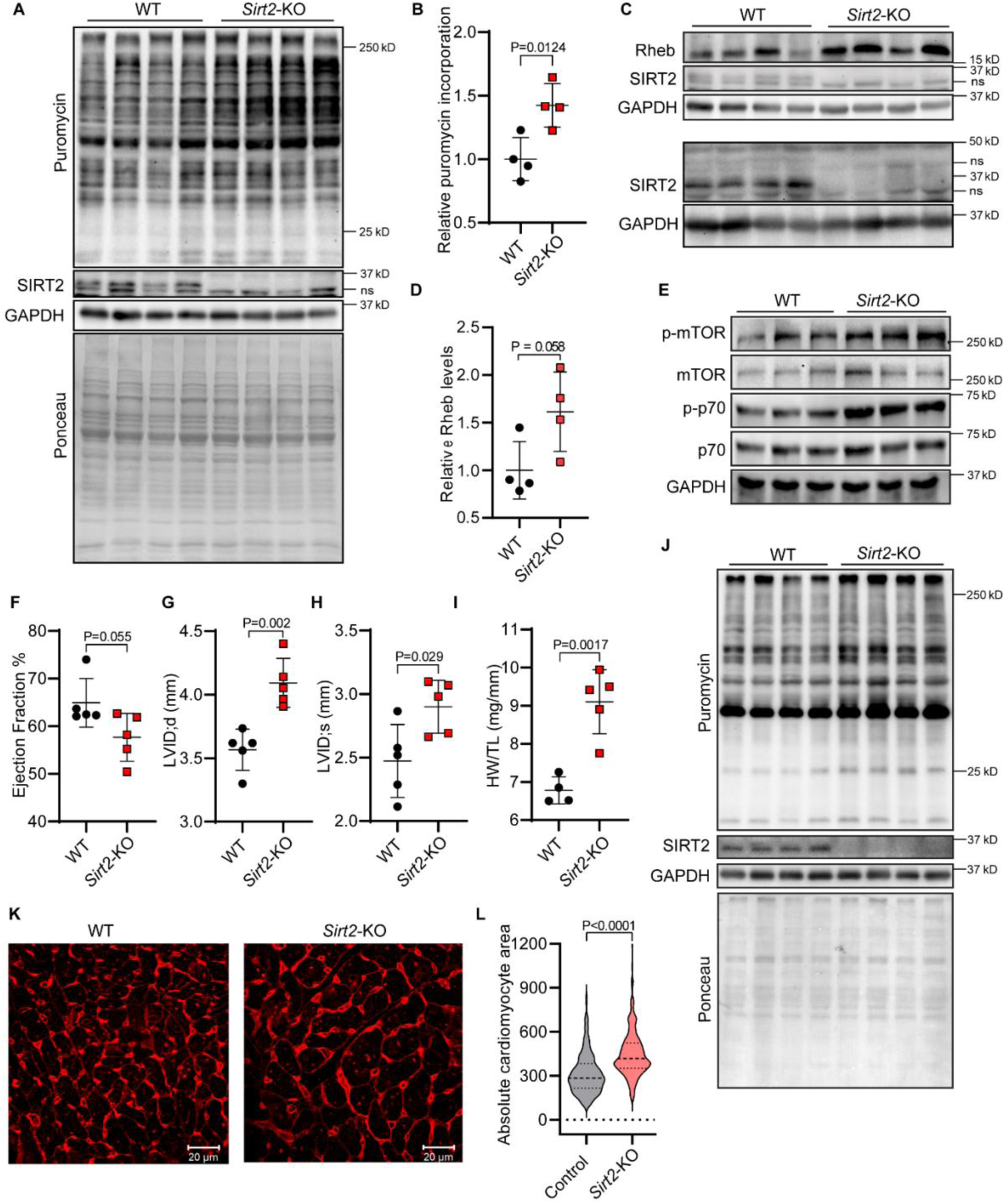
*Sirt2*-KO mice hearts exhibit hyperactivated mTOR kinase activity concomitant with an increased rate of protein synthesis. **(A)** Representative western blot images of SUnSET analysis depicting changes in protein synthesis rate in the heart tissues of 4-month-old *Sirt2*-KO and WT mice. **(B)** Quantification of puromycin incorporation depicted in Figure 6A. The results are expressed as the fold change relative to WT controls. P values shown are from unpaired, 2-tailed Student’s t-test. Data are presented as mean ± s.d, n = 4 mice per group. **(C)** Representative western blot images depicting changes in Rheb protein levels in heart tissues of 4 months *Sirt2*-KO and WT mice. Separate SDS-PAGE was run for increased time and immunoblotted for SIRT2 to show distinct SIRT2 bands via better resolution. **(D)** Quantification of Rheb levels depicted in Figure 6C. The results are expressed as the fold change relative to WT controls. P values shown are from unpaired, 2-tailed Student’s t-test. Data are presented as mean ± s.d, n = 4 mice per group. **(E)** Representative western blot images of analysis of mTOR signaling in the heart tissues of 4-month *Sirt2*-KO and WT mice. **(F)** A representative graph of echocardiographic analysis showing changes in the ejection fraction of 9-month-old *Sirt2*-KO mice when compared with age-matched wild-type mice controls. P values shown are from unpaired, 2-tailed Student’s t-test. Data are presented as mean ± s.d, n = 5 mice per group. **(G)** A representative graph of echocardiographic analysis showing changes in Left ventricular internal diameter, diastolic (LVID; d) of 9-month-old *Sirt2*-KO mice when compared with age-matched controls. P values shown are from unpaired, 2-tailed Student’s t-test. Data are presented as mean ± s.d, n = 5 mice per group. **(H)** Representative graph of echocardiographic analysis showing changes in Left ventricular internal diameter, systolic (LVID; s) of 9-month-old *Sirt2*-KO mice when compared with age-matched controls. P values shown are from unpaired, 2-tailed Student’s t-test. Data are presented as mean ± s.d, n = 5 mice per group. **(I)** Scatterplot representing the heart weight of 9-month-old *Sirt2*-KO and WT mice; the heart weight is normalized with the respective mice’s Tibia length. The results are expressed as the fold change relative to WT controls. P values shown are from unpaired, 2-tailed Student’s t-test. Data are presented as mean ± s.d, n = 4 mice per group. **(J)** Representative images of western blotting SUnSET analysis depicting changes in protein synthesis rate in heart tissues of 9-month-old Sirt2-KO mice and age-matched wild-type control, n=5 mice per group. **(K)** Representative confocal images of WGA-stained cardiac tissue sections showing the cross-sectional area of cardiac cells in WT and *Sirt2*-KO mice at 9 months of age. Scale bar = 20 μm. **(L)** Violin plot showing quantification of WGA images represented in Figure 6(K); P values shown are from unpaired, two-tailed t-test. P values shown are from the Mann-Whitney test. data are shown as median with X and Y percentile. n=5.

### Cardiac-specific SIRT2 overexpression causes a reduction in mTOR activity and protein synthesis in mice hearts

Next, to understand if cardiac-specific overexpression of SIRT2 can reduce protein synthesis in these mice hearts, we developed cardiac-specific SIRT2 overexpressing (csSIRT2-tg) mice **[Figure 7A]**. We confirmed cardiac-specific overexpression through western blotting **[Figure 7B]**. We tested for the protein synthesis levels in these young mice heart tissue by SUnSET assay. We found that protein synthesis was significantly downregulated in the heart tissue of csSIRT2-tg mice compared to flox controls **[Figure 7C, D]**. We observed a reduction in mTOR phosphorylation in the heart tissue of csSIRT2-tg mice compared to flox controls **[Figure 7E]**. These findings suggest that csSIRT2-tg mice show reduced protein synthesis levels and decreased mTOR activation.

**Figure 7.**
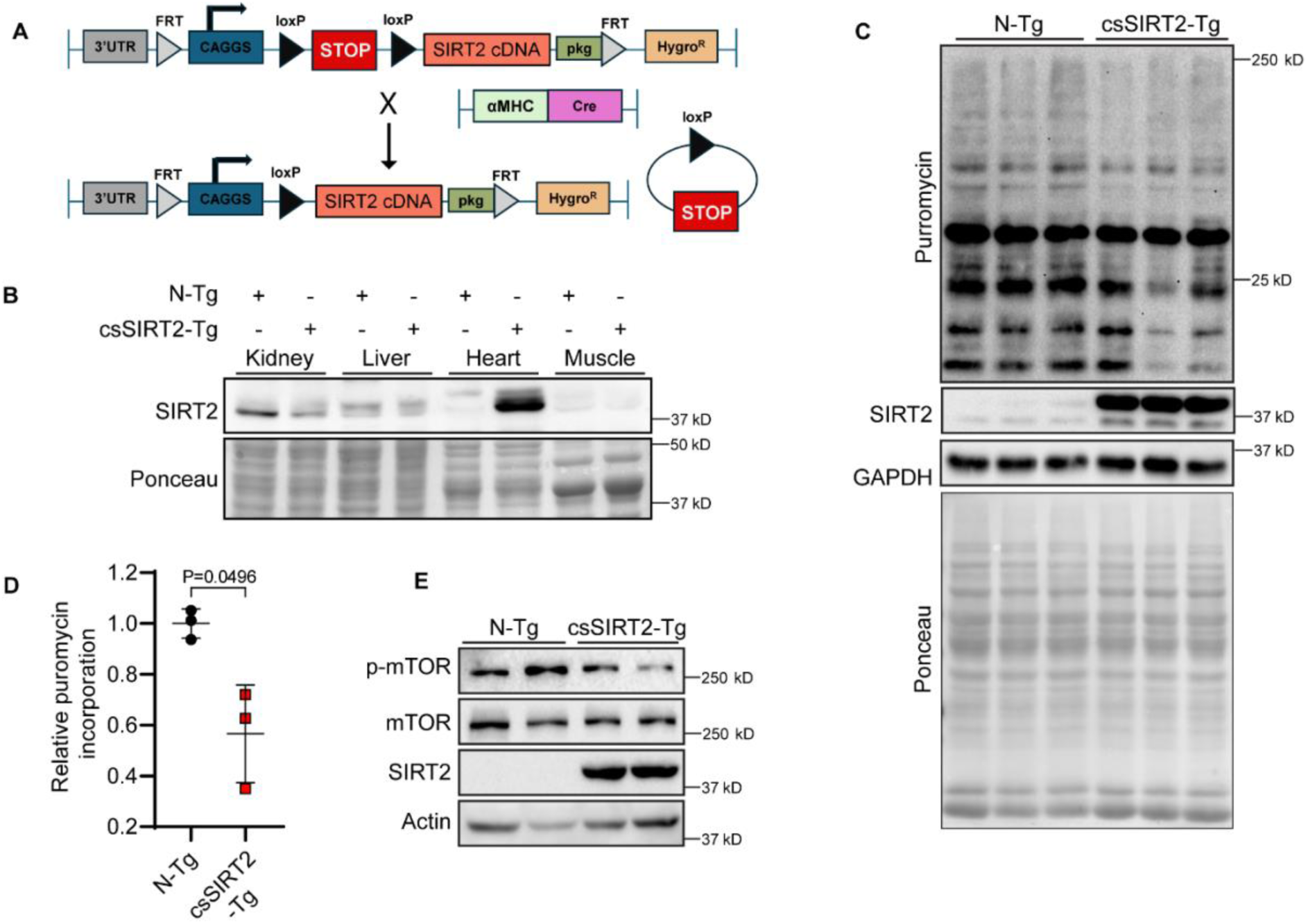
csSIRT2-tg mice hearts display reduced mTOR kinase activity concomitant with a reduced rate of protein synthesis. **(A)** Schematic diagram representing the generation of the cardiac-specific SIRT2-transgenic (csSIRT2-Tg) mice line. **(B)** Representative images of western blotting analysis in different organs of csSIRT2-Tg and control mice showing cardiac-specific overexpression of SIRT2 in csSIRT2-Tg mice heart tissues. **(C)** Representative images of western blotting SUnSET analysis depicting changes in protein synthesis rate in the heart tissues of Non-Transgenic(N-Tg) controls and csSIRT2-Tg mice. **(D)** Scatterplot representing the quantification of puromycin incorporation depicted in Figure 7C. The results are expressed as the fold change relative to non-transgenic controls. P values shown are from unpaired, 2-tailed Student’s t-test. Data are presented as mean ± s.d, n = 3 mice per group. **(E)** Representative western blotting images depicting changes in mTOR activation (mTOR phosphorylation) in heart tissues of Non-Transgenic(N-Tg) controls and csSIRT2-Tg mice.

## Discussion

Our work demonstrates a new role of SIRT2 in regulating protein synthesis. We uncovered Rheb as a novel substrate of SIRT2 and demonstrated a unique regulation of Rheb protein level in cells by its post-translational modifications. Specifically, SIRT2-mediated deacetylation of Rheb on K151 promotes Rheb ubiquitination and proteasomal degradation; Rheb is a crucial activator of mTORC1. Thus, this regulation, in turn, leads to the downregulation of mTOR activity and, thus, protein synthesis.

SIRT2’s effect on protein translation also holds true in mouse models and regulates cardiac hypertrophy in mice. Multiple reports have demonstrated that SIRT2 deficiency in the heart leads to age-associated and neurohormonal-induced cardiac hypertrophy (Tang et al., 2017). We observed increased protein synthesis rates in young *Sirt2*-KO mice heart tissues, increased Rheb levels, and upregulated mTOR activity. We believe that a continuous load of increased protein synthesis in the hearts of these mice contributes to the development of cardiac hypertrophy during aging. Furthermore, studies have shown that Rheb levels are upregulated in pressure overload and aging-induced hypertrophic heart (Blackwood, Hofmann et al., 2019, Ravi et al., 2019), suggesting that regulation of Rheb levels is crucial for cardiac health. We generated a cardiac-specific SIRT2 overexpressing mice model. We found that the heart tissues of these mice heart show a decreased rate of protein synthesis along with decreased mTOR kinase activity, suggesting that csSIRT2-Tg mice protect against the development of cardiac hypertrophy. Moreover, a recent study from Tang et al. (Tang et al., 2017) suggests that csSIRT2-Tg mice are protective against Ang II-induced cardiac hypertrophy. Our findings suggest that SIRT2-mediated check on protein synthesis might play a protective role in csSIRT2-Tg mice against cardiac hypertrophy.

While we focus on cardiac hypertrophy, the SIRT2-Rheb-mTORC1 regulation axis may also be relevant for other human diseases. Deregulated mTOR is strongly associated with various pathologies like arthritis, cancers, neurological disorders, osteoporosis, and insulin resistance (Benjamin, Colombi et al., 2011). SIRT2 has also been connected to several human diseases. For example, SIRT2 is reported to improve insulin sensitivity, and inhibition of SIRT2 displays enhanced progression of diabetic osteoarthritis. SIRT2 also has a disease-promoting role in neurological disorders, positively associated with the progression and development of multiple neurological disorders (Zhu, Dong et al., 2022). We suspect that the SIRT2-Rheb-mTORC1 regulatory pathway may explain the diverse role of SIRT2 in various human diseases.

Sirtuin activities are thought to be regulated by calorie restriction and nutrients availability. This has been demonstrated for SIRT1 and, more recently, for SIRT2 (Guarente, 2013, Wang, Nguyen et al., 2007, Zullo, Simone et al., 2018). It has been reported that SIRT2 levels and deacetylase activity are upregulated under amino acid starvation (Sun, Xiong et al., Zullo et al., 2018). Recently, we reported that SIRT2 deacetylase activity is required to inhibit lipogenesis when cells undergo nutrient stress, mainly amino acid deficiency (Karim, Teng et al., 2024). The current study also observed that complete cell starvation increases SIRT2 levels and activity concomitant with decreased global protein synthesis. These studies suggest that SIRT2 is a crucial nutrient-responsive protein that targets multiple cellular processes in response to nutrient availability.

Moreover, it has been reported that Rheb undergoes K48-linked polyubiquitination under low glucose availability, leading to the inactivation of mTORC1 (Li, Huang et al., 2024), suggesting the essentiality of regulating Rheb levels and activity during glucose starvation, and it may be possible that increased SIRT2 levels might contribute to regulating Rheb levels during nutrients deprivation. Additionally, we also acknowledge that SIRT2 may have other substrates due to which translation is suppressed. Interestingly enough, in another manuscript under submission by Prof. Hening Lin’s lab, SIRT2 is shown to deacetylate 4EBP1 leading to its stabilization, thereby suppressing translation by inactivating mTORC1. This further strengthens the current study by showing that SIRT2 modulates protein synthesis by deacetylating different substrates.

In Summary, our study highlights the intricacy of protein synthesis regulation, a process that holds a controlling influence on cell health and survival. The finding that SIRT2, through Rheb and mTORC1, regulates protein synthesis and cardiac hypertrophy, may also help to uncover new targets to treat cardiovascular diseases.

## Materials and methods

### Cell culture, treatment, and transfection

HeLa cells were grown in a high glucose-containing DMEM medium with 10% fetal bovine serum supplement along with antibiotic-antimycotic. After seeding, cells were subjected to mix humified incubator at 37°C, 5% CO2. The starvation experiment was performed by starving cells for 4hr using EBSS (Earle’s balanced salt solution). Transfection was performed in cells when they were at 70-80% confluency. Lipofectamine®3000 reagent was used for the plasmid transfection, whereas Lipofectamine RNAiMAX was used for the transfection of siRNAs, all these transfections were performed according to the manufacturer’s protocol, and the cells harvested at 48 hours post-transfection in case of plasmid-mediated overexpression and at 72 hours for siRNAs mediated gene knockdown. AGK2 treatment (10μM) was used for the inhibition of SIRT2 activity. Protein synthesis rate was measured through the Surface Sensing of Translation (SUnSET) assay as described in our previously published study (Ravi, Jain et al., 2020). Puromycin (1μM) was added to the complete DMEM media 30 minutes before cell harvesting.

### Animal Studies

All animal studies were carried out after getting approval from the Institutional Animal Ethics Committee of IISc, Bengaluru, India. The animals were given a normal chow diet and were considerably maintained in cages provided with good ventilation. A 12-hour light/dark cycle was followed for these animals throughout the study. For the *in vivo* SUnSET assay, mice were injected with puromycin. Puromycin was given at a dose of 40nmol/g of body weight via an intraperitoneal injection. Post 30 minutes of puromycin administration, the mice were sacrificed, and tissue harvesting was performed. Harvested tissues were snap-frozen and stored at -80°C until further use. 4-months and 9-months age-matched wild-type control, and SIRT2 Knockout (*Sirt2*-Ko) mice were used for the SUnSET assay. Two-month-old control and csSIRT2-Tg were administered with puromycin to evaluate the protein synthesis rate and mTOR kinase activity. Change in mTOR activation was also evaluated in the same age-grouped csSIRT2-Tg mice. Cardiac functions of *Sirt2-*KO mice at 4,9,12 months and age-matched control mice were examined via Visual Sonics high-frequency ultrasound system as mentioned previously (Sarikhani, Maity et al., 2018b).

### Cell and tissue lysate preparation

Cells were harvested from their respective culture dishes by using a cell scraper after adding required volume of RIPA lysis buffer (20 mM Tris–HCl pH 7.5, 1 mM EDTA, 150 mM NaCl, 1 mM EGTA, 1%NP-40 1%, 1% sodium deoxycholate, 1 mM sodium orthovanadate, 2.5 mM sodium pyrophosphate, 1 mM PMSF and 1X protease inhibitor cocktail). The same buffer was used to lyse tissue samples. Prior to the lysis, Tissue samples were frozen in liquid N_2_ and grinded by using mechanical approach. Centrifugation at 13000 rpm, 10 minutes at 4°C was performed to clear out the lysate, and the supernatant was collected and stored at -80°C to ensure long-term stability.

### Site-directed mutagenesis

Rheb mutants, K151Q, and K151R were constructed by using site-directed mutagenesis of pRK5 HA GST Rheb1 plasmid. We have used QuikChange II Site-Directed Mutagenesis Kit (Agilent Technologies) and performed SDM according to the manufacturer’s protocol. These mutants contain a single point mutation of indicated Lysine (K) residues, where Lysine has been replaced by either Arginine (R) or Glutamine (Q). SIRT2 catalytic mutant N168A was also generated using the same kit as mentioned above, and all these mutants were confirmed by sequencing analysis.

### Electrophoresis and Immunoblotting

Quantification of protein lysates was done by using Bradford assay (Bio-Rad). The protein samples were normalized by using an equal amount of protein. 2x Laemmle sample buffer was added to the lysates along with 5% beta-mercaptoethanol, and the mixture was boiled at 95°C for 5 minutes. Samples were subjected for electrophoresis on 10% SDS-PAGE gels. After the adequate time of electrophoresis, proteins were transferred onto 0.45 45μm PVDF membranes by using fast or slow wet transfer. After the completion of transfer the membrane was then blocked for 1hr at RT with 5% skim milk prepare in 1X TBST (Tris-Buffered Saline supplemented with 0.05% Tween 20). Specific antibodies diluted either in 5% BSA in TBST or 5% Skim milk in TBST were added to the membranes followed by overnight incubation at 4°C. The membrane-bound primary antibodies were then recognized by using HRP-conjugated secondary antibodies diluted in 1% skim milk for 1 hr at RT. Washings in the subsequent steps were done by using TBST. Chemiluminescence signal was detected with the use of ECL reagents, and the images were obtained with the help of a Chemiluminescence imager.

### Immunofluorescence microscopy

To perform immunofluorescence microscopy-based studies, the cells were first seeded on sterile glass-made coverslips in a 12-well plate. After subsequent transfections and treatment, at the experimentally decided time, cells were washed thrice with 1X PBS. After completely draining the PBS out the cells were fixed by using 4% paraformaldehyde (PFA) for 15 minutes at RT. Upon fixation, the extra PFA was removed by giving PBS washes, and the fixed cells were permeabilized and blocked in a single step with 0.1%Saponin-5% BSA solution in PBS for 1hr at RT. After the blocking, specific antibodies prepared in 1% BSA-saponin in PBS solution were added to the cells. Cells were then incubated overnight at 4°C. After the completion of Respective secondary antibodies conjugated with Alexa Fluor 488 and/or Alexa Fluor 594 prepared in 1% BSA-saponin in PBS along with DAPI were added to the cells for 1hr. the cells were washed three times with 1X PBS and mounted on a clean ethanol-wiped glass slide by using Fluoromount G. Images were obtained with a Zeiss LSM 710 or 880 confocal microscope.

### Immunoprecipitation assay

To check protein-protein interaction, HeLa cells were lysed using cell lysis buffer (20 mM Tris–HCl pH 7.5, 1 mM EDTA, 150 mM NaCl, 1 mM EGTA, 1% Triton X100, 1 mM sodium orthovanadate, 2.5 mM sodium pyrophosphate, 1 mM PMSF and 1X protease inhibitor cocktail). A total of 750-1000μg protein was incubated with the 2μg of specific antibody. This antibody-protein mixture was incubated overnight at 4°C, 5 rpm in a rotator. Upon overnight incubation, immunoprecipitated complexes were pulled down from the mixture by using magnetic protein G Sepharose beads (Bio-Rad). After subsequent washing with the PBS, these beads were resuspended in 2X: Laemmli buffer and the mixture was subjected to heat-mediated immunoprecipitants dissociation at 95°C for 5 minutes; further 5% Beta-mercaptoethanol was added, and the mixture was again incubated at 95°C for 2 minutes, and Proteins was detected using western blotting. For Rheb-SIRT2 interaction, endogenous Rheb was pulled down using a Rheb-specific antibody. Western blotting analysis was performed and Rheb-SIRT2 interaction was confirmed using Rheb and SIRT2 specific antibodies.

### Cap-dependent translation assay

pcDNA3-EMCV bicis plasmid was used to evaluate Cap-dependent translation in cells under different treatment conditions. Briefly, this plasmid contains R Luc under a 5’ caped site (representing Cap-dependent translation) and F Luc under an EMCV IRES sequence (used as internal control). This bicis plasmid was transfected in HeLa cells under different treatment conditions. Lipofectamine 3000 was used as a transfection reagent to enhance transfection efficiency. Luciferase readings were measured using a dual luciferase reagent kit (Promega), as per the manufacturer’s protocol.

### Protein Expression and Purification

Human Ras homolog enriched in brain (Rheb) 1-169 aa and its mutants K151 and K151Q were expressed using GST-tagged pGEX 4T vector in Rossetta BL21 cells. Expression was induced using 0.5 mM IPTG at 0.6 OD600 for 5 hours at 37 °C and cells were pelleted at 4500 rpm. ^15^N labelled and unlabelled hRheb and mutants were expressed in similar manner.

Sonication/lysis buffer 50 mM Tris, pH 7.5, 150 mM NaCl, 1 mM MgCl2 was used for resuspension of pellets and cell lysis was performed using sonication. Lysed cells were subjected to centrifuge at 13000 rpm for one hour. Supernatant was loaded on pre-packed GST column. Bound GST tagged Rheb or the mutants were eluted in 50 mM Tris, 100 mM NaCl, 10 mM reduced Glutathione, pH 8.0. GST tag was removed using thrombin by incubating for 16 hours at 4-8 °C. After incubation period, 0.1-1 mM PMSF was used to inhibit thrombin followed by sample was subjected to gel filtration (S75 column) to separate GST and Rheb (and mutants). During gel filtration, Rheb or mutants were eluted in 50 mM Phosphate, pH 7.5, 100 mM NaCl 5 mM MgCl2.

### NMR experiments

2D ^1^H-^15^N HSQC NMR spectra of Rheb without GTP analog (GppNHp) and with GTP analog were recorded on Bruker 700 MHz spectrometer at 298 K. 180 μM Rheb was prepared in 50 mM Phosphate, pH 7.5, 100 mM NaCl 5 mM MgCl_2_ and same buffer was used for GTP analog (GppNHp) solution. Titration experiments were performed for Rheb WT, Rheb K151Q and Rheb K151R with GppNHp at five different ratio (1 : 0.25, 1 : 0.5, 1 : 1, 1: 2 and 1 : 5). All the samples contained 10% D2O for the spectrometer deuterium lock. All NMR results were processed using Bruker TOPSPIN 3.1 and data were analyzed using NMRFM-SPARKY (Delaglio, Grzesiek et al. 1995, Lee, Tonelli et al. 2015). Resonance assignment of few residues were taken from a previously published assignment from Biological Magnetic Resonance Data Bank (BMRB) entry 15202 (Schwarten, Berghaus et al., 2007).

### Pulling simulation (Umbrella sampling)

To examine binding of GDP with WT Rheb, Rheb K151Aly, Rheb K151K and Rheb K151Q pulling force simulation was performed (Lemkul & Bevan, 2010). Molecular topology of all four systems were generated using Charmm36 forcefield parameters included in GROMACS package. Box dimension was set for 6.560 x 4.362 x 12 and center of mass is placed at 3.280, 2.181, 2.4775 in this box. Energy minimization and equilibration (NPT) were performed similarly as mentioned MDS. Over a period of 500 ps, GDP was pulled away from all four Rheb systems, and during this 500 ps, 500 configurations (also called reaction coordinates) were generated. Results were analyzed using Grace plotting tool.

### Rheb Acetylation assay

Flag or HA-tagged Rheb plasmids were transfected into 293T control and SIRT2 knockdown cells using PEI reagent. An empty plasmid with a C terminal flag tag was used as a negative control. Cells were collected the day after the transfection and were washed with PBS prior to collecting via centrifugation of 3000 rpm for 5 minutes at 4 °C. 1 ml of 1% NP40 lysis buffer with final pH 7.4 (150 mM Tris-HCl, pH 8, 150 mM NaCl, 10% glycerol, 1% NP40) was used to lyse each sample at 4 °C for 1 hour. After spinning down the samples at 13000 rpm for 10 minutes at 4 °C, the supernatant was collected and normalized after using the Bradford reagent to determine the protein concentration. 40ul of each normalized sample was collected as input and was boiled at 95 °C for 5 minutes after 8 ul of 6 times loading dye was added to each input sample. The rest of the normalized samples were incubated with 20 ul of HA, Flag, or Acetyl Lysine beads at 4 °C overnight. The affinity beads were then washed with IP washing buffer (25 mM Tris, pH 8.0, 150 mM NaCl, 0.2% NP40) for 3 times and were dried with a 1 ml syringe. Depending on the experiment, 50 ul of 1x loading dye or 35 ul of 2x loading dye were added to the dried beads, and each sample was boiled at 95 °C for 5 minutes. Standard western blot analysis was used to analyze the samples.

### Molecular Simulation studies

All-Atom Simulations: A series of NPT simulations were performed on four systems, each with the same initial orientation of Rheb protein placed on the membrane. For Rheb, structure deposited as RCSB PDB: 1XTS was used (Yu, Li et al., 2005). The systems differed mainly in protein mutations i.e. (a) Wildtype (WT), (b) K151A (mutant), (c) K151Q (Acetylated mimic) and (d) K151R (Deacetylated mimic). Lipid membranes were generated using CHARMM-GUI online server (Jo, Kim et al., 2008). The membrane composed of identical 80:20 mixture of POPC, and POPS for all the four systems. The bilayer PHD system was energy minimized in vaccum. Further, the systems were solvated with TIP3 water molecules (Mark & Nilsson, 2001) and Potassium and Chloride ions were added to neutralize each system and bring the salt concentration to 150 mM. Each system is simulated using GROMACS (Pronk, Páll et al., 2013) with CHARMM36m protein force field (Huang, Rauscher et al., 2017) and the CHARMM36 lipid force field (Klauda, Venable et al., 2010) with a 2-fs time step and visualized using VMD (Humphrey, Dalke et al., 1996). Simulations were performed for 500 ns at T=303K and 1 atm pressure. Systems were minimized using steepest descent algorithm followed by the equilibration of 5-10ns. Verlet Cutoff-scheme was used. Frequency to update the neighbor list (nstlist) was taken as 20. Periodic boundary conditions were used in all directions i.e. xyz was used for pbc. Temperature was coupled using Nose – Hoover thermostat (Evans & Holian, 1985). Pressure coupling was performed using Parrinello - Rahman barostat (Parrinello & Rahman, 1981) using semi-isotropic coupling scheme. Non-bonded forces were calculated with a 12 Å cutoff (10 Å switching distance). Long-range electrostatic forces were calculated every other time step using the particle mesh Ewald method (Essmann, Perera et al., 1995). The system is maintained at a temperature of 303 K. The choice of the initial systems was guided by inputs from Orientations of proteins in membranes (OPM) database (Lomize, Lomize et al., 2006) and from a published work by Gorfe group (Prakash & Gorfe, 2022). For carrying out the z-analysis for membrane association, in-house tcl script was used.

### Mass spectrometry sample preparation

Rheb was immunoprecipitated from control, and AGK-2-treated Hela cells, and the samples were run in SDS-PAGE until the entire volume of samples was stacked in SDS-Polyacrylamide gel. Gel was cut at the stacked area using a sharp, sterile blade for further processing. The gel piece was treated with 5 mM TCEP and alkylated with 50 mM iodoacetamide. Shrink the gel band with Acetonitrile and air-dry for a few minutes at room temperature, then digested with trypsin (1:50, trypsin/lysate ratio), for 16 h at 37 °C. Digests were speed vac for 1 hour, and the pellet was dissolved in Buffer A (5% acetonitrile, 0.1% formic acid) and then cleaned using a C18 silica cartridge and dried using a speed vac. The dried pellet was resuspended in buffer A (5% acetonitrile, 0.1% formic acid). Analysis was performed using an EASY-nLC 1000 system (Thermo Fisher Scientific) coupled to a QExactive mass spectrometer (Thermo Fisher Scientific) equipped with a nanoelectrospray ion source. 1.0 µg of the peptide mixture was resolved using a 15 cm PicoFrit column (360µm outer diameter, 75µm inner diameter, 10µm tip) filled with 1.9 µm of C18-resin (Dr Maeisch, Germany). The peptides were loaded with buffer A and eluted with a 0–40% gradient of buffer B (95% acetonitrile, 0.1% formic acid) at a flow rate of 300 nl/min for 45 min. MS data was acquired using a data-dependent top10 method dynamically choosing the most abundant precursor ions from the survey scan. RAW files obtained in this study were analyzed using Proteome Discoverer against the sequence given. For Sequest search, the precursor and fragment mass tolerances were set at 10 ppm and 0.5 Da, respectively. The protease used to generate peptides, i.e., enzyme specificity, was set for trypsin/P (cleavage at the C terminus of “K/R: unless followed by “P”) along with maximum missed cleavage value of two. Carbamidomethyl on cysteine as fixed modification and oxidation of methionine, N-terminal acetylation (K), and lysin Ubamide were considered variable modifications for database search. Both peptide spectrum match and protein false discovery rate were set to 0.01 FDR.

### Histology

Heart tissue samples were collected in 10% neutral buffered formalin soon after euthanasia. After fixing samples (seventy-two hours incubation in formalin), formalin was withdrawn from the samples by putting the samples overnight under a running water tap (minimum 9 hours.). When the formalin was eliminated, samples were subjected to dehydration in alcohol and then xylene, and later covered with paraffin wax by using an automated tissue processor (Leica TP1020, Germany). Then, Samples were embedded using paraffin wax to make blocks to ensure easy sectioning (Leica EG 1150H, Germany). These Blocks were then solidified using a cooling machine (Leica EG 1150C, Germany). Sections were cut (5µm thickness) and put on glass slides with the help of a microtome (Leica RM2245, Germany). Wheat Germ Agglutinin (WGA) (5µg/ml) stain was used to evaluate the cross-sectional area of cardiomyocytes.

### Click Chemistry

Flag-Rheb plasmids were transfected into HEK293T control and SIRT2 knockdown cells using PEI. 24 hours later, half of the samples were treated with 50 μM Alk14 for 6 hours before collecting. The samples that were not treated with Alk14 were served as negative controls. Samples were collected via centrifugation of 3000 rpm for 5 minutes at 4 °C. 1 ml of 1% NP40 lysis buffer with final pH 7.4 (150 mM Tris-HCl, pH 8, 150 mM NaCl, 10% glycerol, 1% NP40) was used to lyse each sample at 4 °C for 1 hour. After spinning down the samples with 13000 rpm for 10 minutes at 4 °C, the supernatant was collected and normalized after using the Bradford reagent to determine the protein concentration. 40 μL of each normalized sample were collected as input and was boiled at 95 °C for 5 minutes after 8 μL of 6 times loading dye was added to each input sample. The rest of the normalized sample were incubated with 20 μL of Flag beads at 4 °C over night. The flag beads were then washed with IP washing buffer (25 mM Tris, pH 8.0, 150 mM NaCl, 0.2% NP40) 3 times and were dried with a 1 ml syringe. Each sample was then added with 24 μL of click chemistry mixture (20 μL IP washing buffer, 1 μL of 40 mM CuSO4, 1 μL of 10 mM TBTA, 1 μL of 2 mM TAMARA-azide, and 1 μL of 40 mM TCEP). The samples were then incubated in the dark for 30 minutes, and 12 μL of 6X SDS loading dye were added to each sample. After boiling at 95 °C for 5 minutes, samples were divided in half, and half of the samples were added with 400 mM hydroxylamine and boiled for another 10 minutes. All samples were then analyzed with SDS-PAGE, and palmitoylation signal was checked using the Rhodamine channel of ChemiDoc Imaging System.

### Quantification and statistical analysis

All statistical analysis and graphical representation of data was performed using Graph-pad Prism version 8.4.2. For pair-wise comparisons, if the data values follow the Gaussian distribution, then a parametric *t-test* with Welch’s correction was used. If the data values were not following Gaussian distribution, then the nonparametric Mann-Whitney test was used. One-way ANOVA and two-way ANOVA were used for comparison between more than two groups. ZEN-Black software was used for confocal image processing and analysis. Image Lab software (BioRad) was used for processing western blot images acquired from the Chemi-Doc machine and ImageJ software was used to quantify western blots.

## Author contributions

N.R.S carried out conceptualization, project administration, and supervision. AJ.S designed and analyzed the experiments and wrote the first draft of the manuscript; Y.Z performed click chemistry, Rheb acetylation, and ubiquitination IP experiments and helped in manuscript editing. AJ.S, D.N, and A.S.P performed western blotting experiments, and A.S.P and AJ.S generated mass-spectrometry data. B.S, AJ.S, and D.N did *Sirt2-KO* mice breeding, maintenance, and harvesting. B.S. performed Echocardiography for SIRT2 KO mice used in this study. D.N. carried out data verification and confocal experiments. B.S and D.K provided csSIRT2-Tg mice used in this study. V.K.S and S.G carried out NMR and pulling force experiments. K.J performed simulation studies to understand the membrane localization of Rheb and Rheb mutants. V.R provided various constructs and reagents used in the study and helped in scientific discussion. S.R helped in manuscript editing and scientific discussions. B.P.A performed SDM to generate Rheb mutants, and T.S.S. performed WGA staining for *Sirt2*-KO mice heart tissues. H.L, A.S, and M.S provided resources, designed experiments, and reviewed the manuscript.

## Resource availability

### Lead contact

Additional information and requests for reagents and resources should be directed to, and will be fulfilled by, the Lead Contact, N. Ravi Sundaresan.

### Materials Availability

All unique/stable reagents generated in this study are available from the Lead Contact with a completed Materials Transfer Agreement.

### Data and code availability

The source data for figures in the paper are available from the Lead Contact on request.

## Acknowledgments

Prof. Saumitra Das (IISc, Bengaluru, India) is acknowledged for providing the EMCV-Bicis construct. We acknowledge the Central Animal Facility, NMR facility, Central Confocal facility, Division of Biological Sciences and the Confocal facility, Department of Microbiology and Cell Biology, Indian Institute of Science, Bangalore, for their services and technical help. N.R.S. acknowledges the funding agency DBT Government of India for providing funds to support this study (BT/PR31212/BRB/10/1759/2019). AJ.S acknowledges MHRD Government of India for providing fellowship to support this study.

## Summary Figure

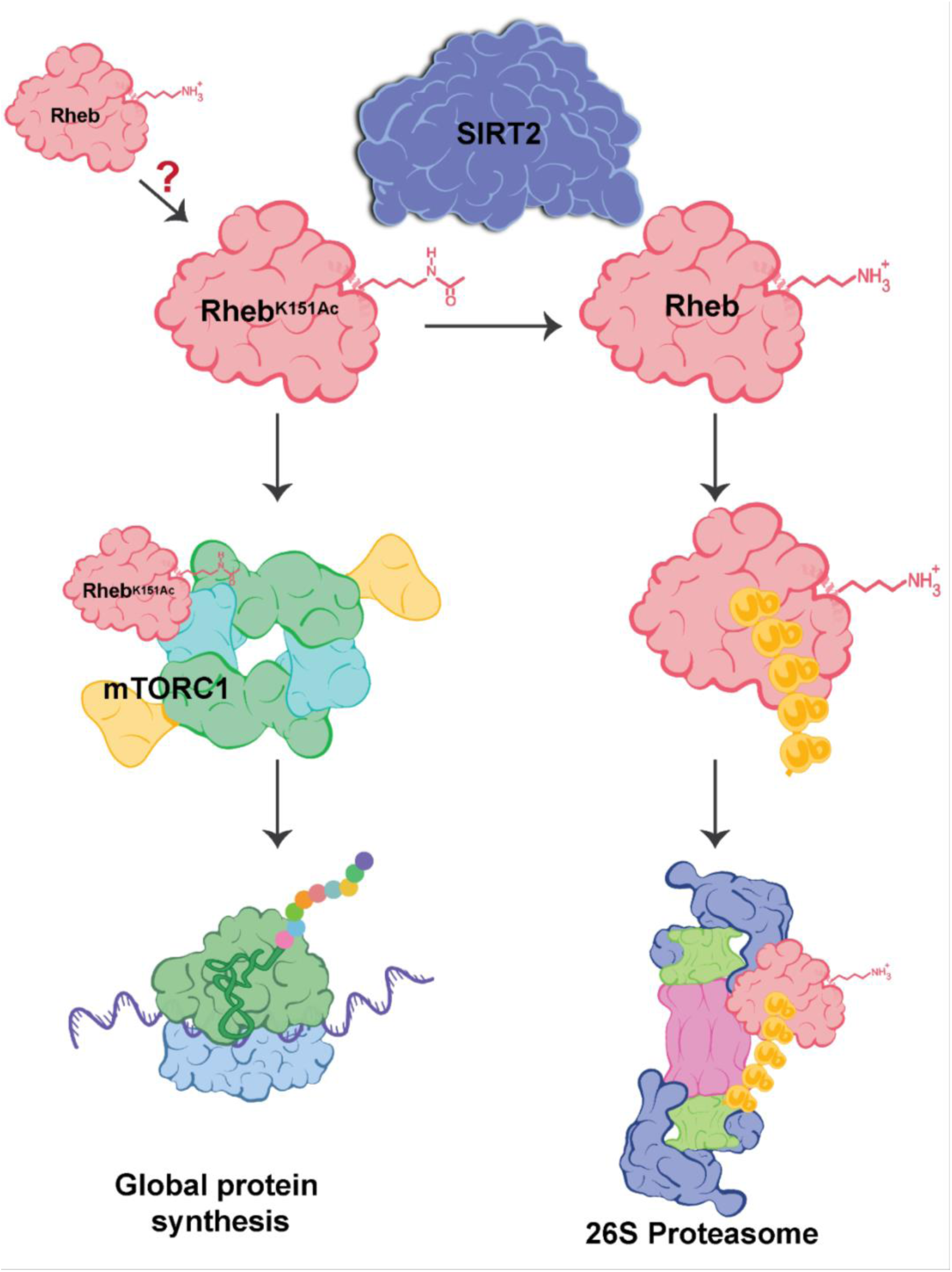

## Supplementary Figures

**Figure S1.**
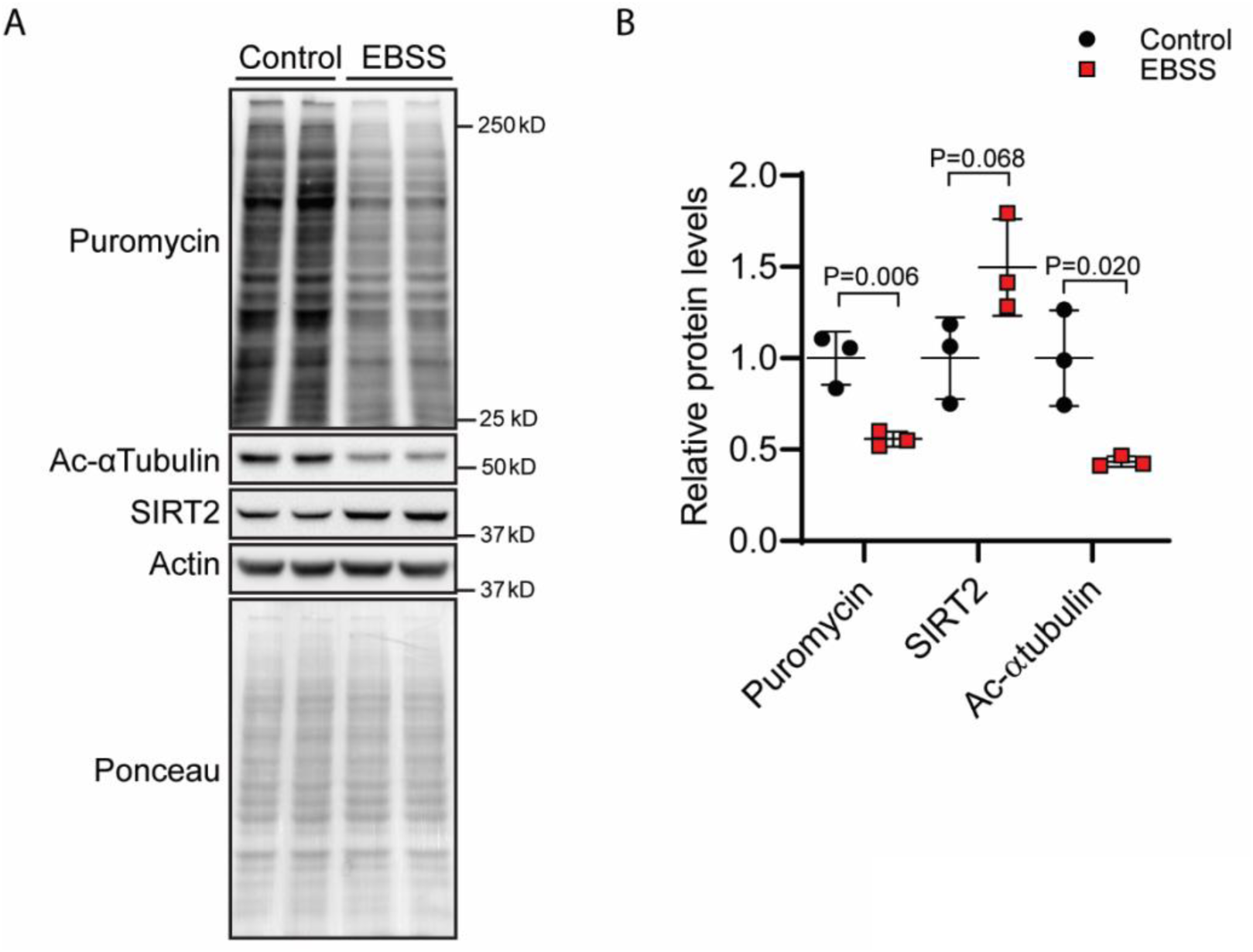
SIRT2 levels and activity are upregulated in nutrients deprived conditions along with decreased protein synthesis rates. **(A)** Representative images of Western blotting analysis indicating the changes in protein synthesis rate (tracked using WB-SUnSET assay) and concomitant changes in the levels of acetyl-tubulin and SIRT2. in HeLa cells treated with high glucose DMEM media supplemented with 10% FBS (CM) and complete starvation using EBSS. **(B)** Scatterplot representing quantification of immunoblots from Figure (S1A). The graph represents relative protein levels of puromycin-incorporated peptides, acetyl-tubulin, and SIRT2. n = 3. Statistical significance was determined by multiple t-tests using the Holm-Sidak method, with alpha = 0.05. Each row was analyzed individually, without assuming a consistent SD. Data are shown as mean ± s.d.

**Figure S2.**
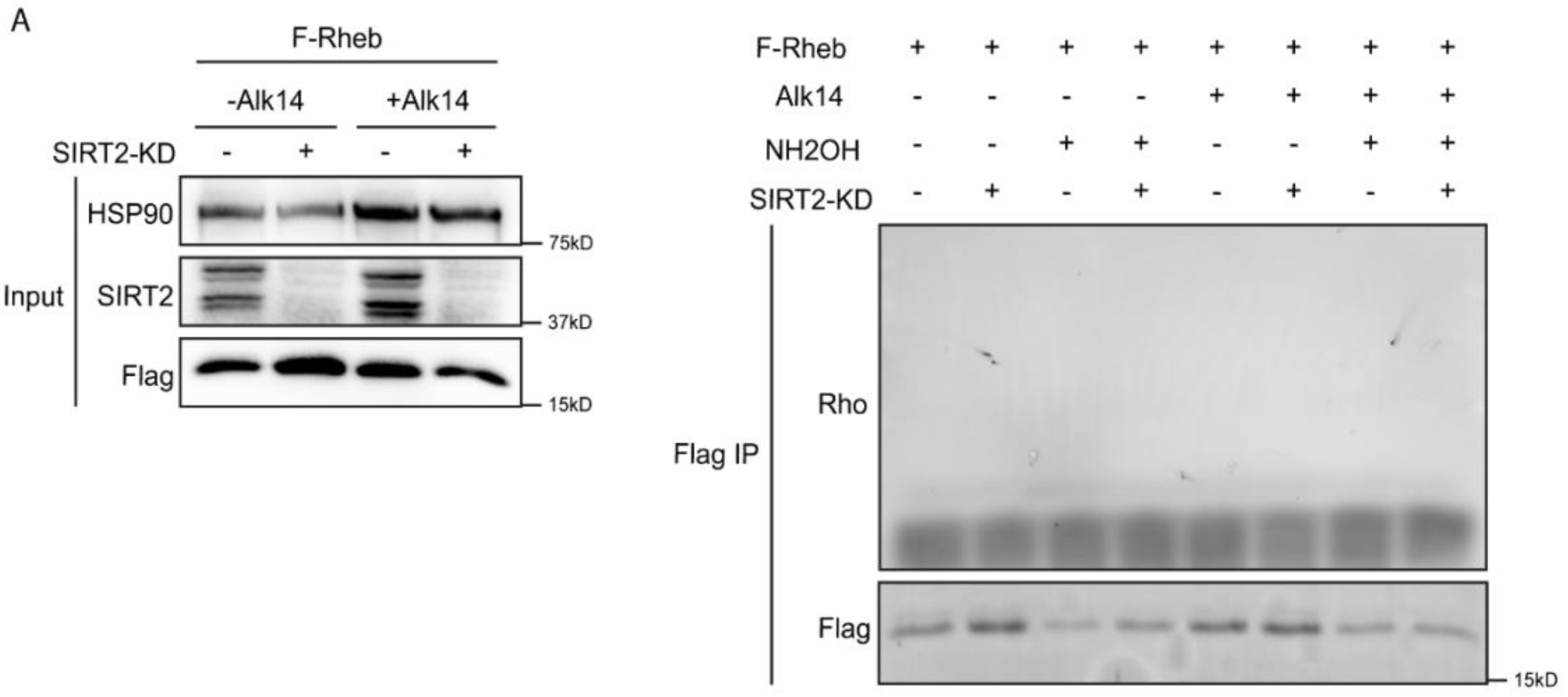
Rheb is not endogenously long-chain fatty acylated under normal conditions. **(A)** Alk14 metabolic labeling was used to detect Rheb palmitoylation. Flag-tagged RheB was overexpressed in control and SIRT2 knockdown HEK 293T cells. Cells were labeled with Alk14, and samples were treated with or without hydroxylamine (which removes cysteine palmitoylation, but not lysine palmitoylation). Click chemistry was performed to conjugate a rhodamine-azide fluorescent label to proteins that are labeled with Alk14. Flag-Rheb was then pulled down and its potential palmitoylation was detected by in-gel fluorescence. The lack of fluorescence signal indicates Rheb is not palmitoylated under the conditions tested.

**Figure S3.**
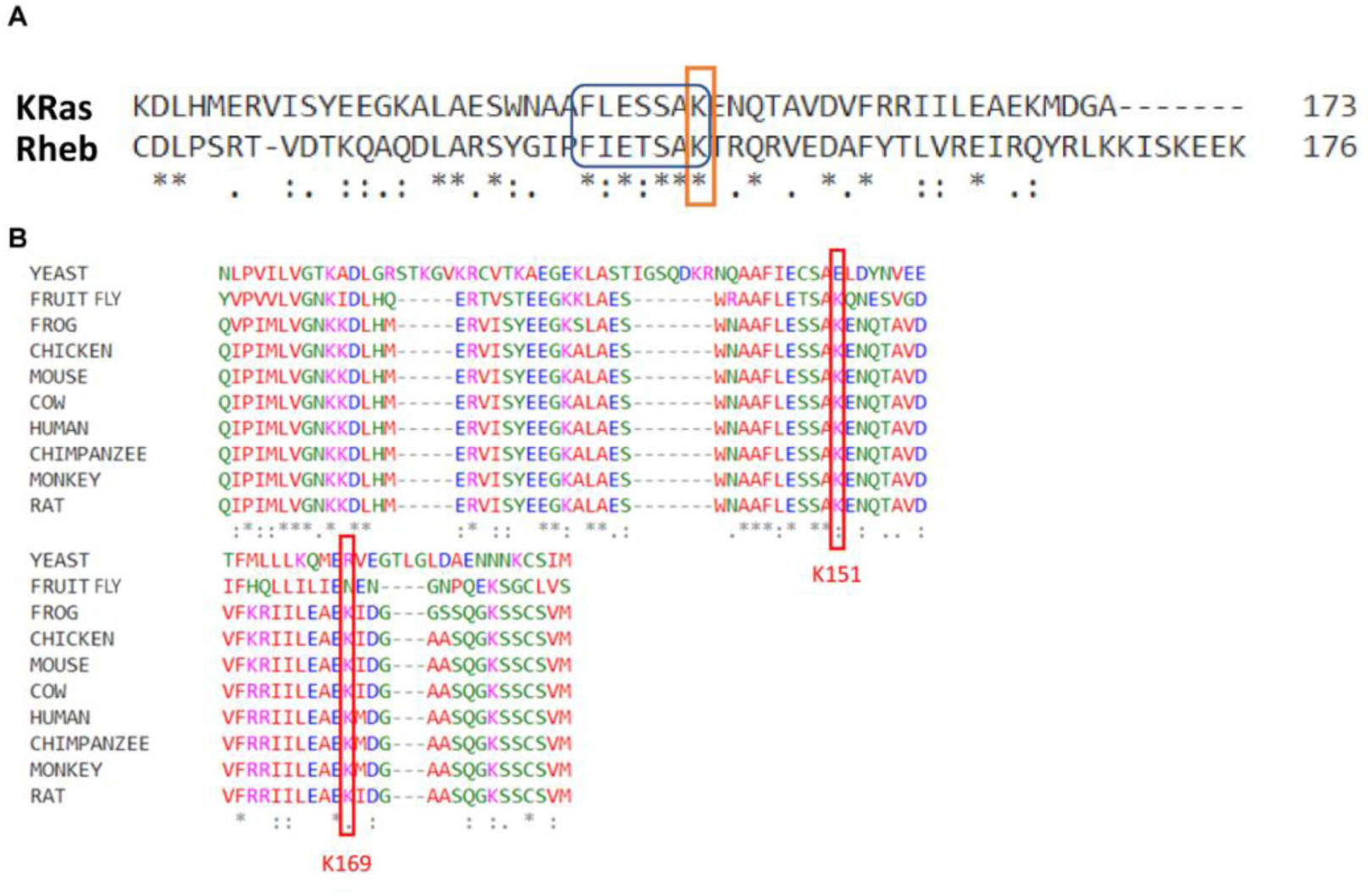
Lysine 151 is a conserved residue across different species; Rheb and KRas share a common sequence stretch for SIRT2-targeted Lysin residue. **(A)** Representative image showing sequence similarity in SIRT2 targeted region of Rheb and KRas, Rheb and KRas share a similar sequence stretch of amino acids for K151 and K147 residues, respectively. **(B)** Sequence alignment data depicting K151 residue in Rheb is conserved across different species.

**Figure S4.**
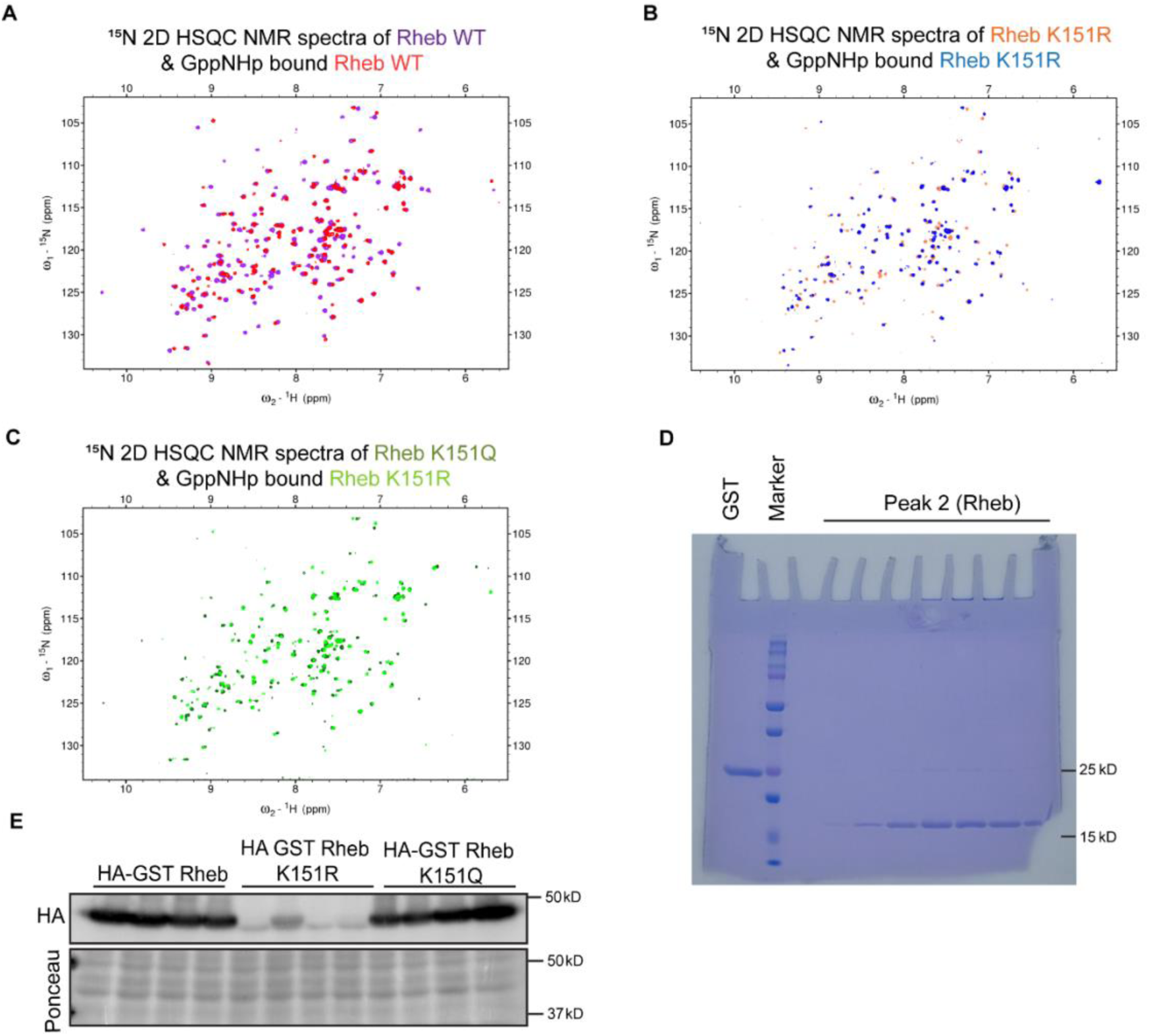
Rheb mutants K151R and K151Q are structurally stable in basal and GppNHp bound form. **(A)** Representative overlaid 2D ^1^H-^15^N HSQC NMR spectra of Rheb WT & GppNHp bound Rheb WT. **(B)** Representative overlaid 2D ^1^H-^15^N HSQC NMR spectra of Rheb K151R & GppNHp bound Rheb K151R. **(C)** Representative overlaid 2D ^1^H-^15^N HSQC NMR spectra of Rheb K151Q & GppNHp bound Rheb K151Q. **(D)** Representative SDS-PAGE gel image showing purified Rheb protein at 15 kD molecular weight. **(E)** Representative images of western blot analysis showing differential expression of HA-GST Rheb, HA-GST Rheb K151R, and HA-GST Rheb K151Q. Ponceau of the same blot is used as a loading control.

**Figure S5.**
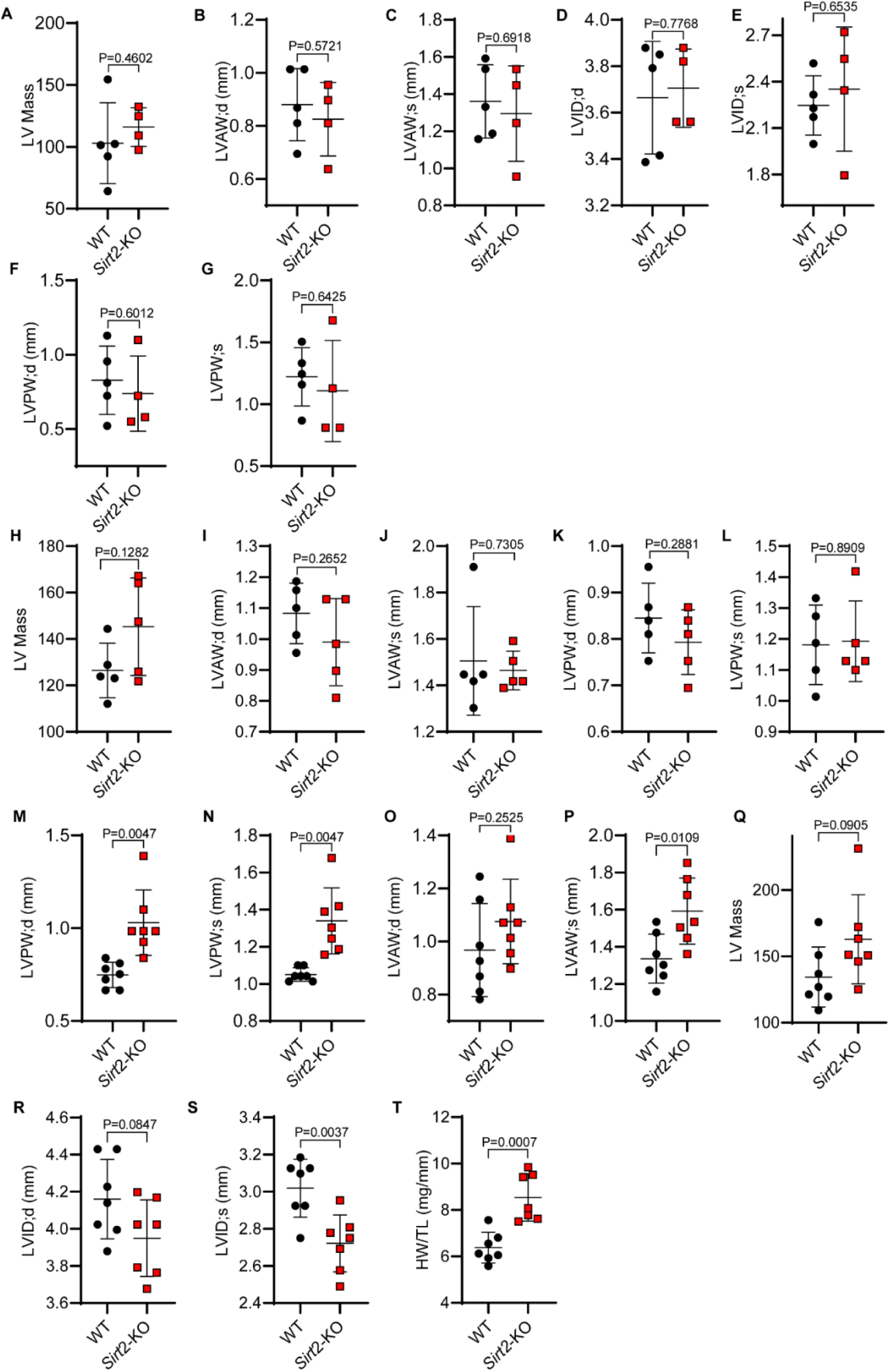
Representative graph of echocardiographic analysis showing changes in the various cardiac parameters of 4,9, and 12-months-old *Sirt2-KO* mice. **A-G**. Representative graph of echocardiographic analysis showing changes in the various cardiac parameters of 4-month-old *Sirt2*-KO mice when compared with age-matched wild-type mice controls. Data are presented as mean ± s.d, n = 4-5 mice per group. P values shown are from unpaired, 2-tailed Student’s t-tests. Data are shown as mean ± s.d, **H-L.** Representative graph of echocardiographic analysis showing changes in the various cardiac parameters of 9-month-old *Sirt2*-KO mice when compared with age-matched wild-type mice controls. Data are presented as mean ± s.d, n = 5 mice per group. P values shown are from unpaired, 2-tailed Student’s t-tests. Data are shown as mean ± s.d, **M-T.** Representative graph of echocardiographic analysis showing changes in the various cardiac parameters of 12-month-old *Sirt2*-KO mice when compared with age-matched wild-type mice controls. Data are presented as mean ± s.d, n = 7 mice per group. P values shown are from unpaired, 2-tailed Student’s t-tests. Data are shown as mean ± s.d,

